# A stratagem for primary root elongation under moderate salt stress in the halophyte *Schrenkiella parvula*

**DOI:** 10.1101/2022.11.03.515116

**Authors:** Keriman Sekerci, Nahoko Higashitani, Rengin Ozgur, Baris Uzilday, Atsushi Higashitani, Ismail Turkan

**Affiliations:** Graduate School of Life Sciences, Tohoku University, Sendai, 980-8577, Japan; Faculty of Science, Department of Biology, Ege University, Bornova, 35100, Izmir, Turkey

**Keywords:** Auxin, Halophyte, Root elongation, Salinity, *Schrenkiella parvula*

## Abstract

Halophytes are salt-tolerant plants that grow in soil or waters of high salinity. *Schrenkiella parvula* is one of the halophyte plants that grow around Tuz (Salt) Lake, TURKEY that can survive at 600 mM NaCl. Intriguingly, *S. parvula* belongs to the same Brassicaceae family as the model plant *Arabidopsis thaliana*, and its genome is 90% homologous to the Arabidopsis genome. Here, we performed proteomic analysis and physiological studies on the roots of *S. parvula* seedlings cultivated under a moderate salt condition at 100 mM NaCl. Surprisingly, under 100 mM NaCl conditions, the primary roots elongated much faster than under NaCl-free conditions, although up to 200 mM those were reduced. On the other hand, iso-osmotic mannitol did not promote primary root elongation, suggesting a specific response to NaCl. Epidermal cell elongation was promoted in the elongation zone, but meristem size and DNA replication were decreased. In addition, root hair formation and lateral root elongation were suppressed at moderate salinity. Compared with *A. thaliana*, the cell death and ROS increase of root tip meristem cells under 100 mM NaCl condition were significantly lower in *S. parvula* seedlings. The size and starch content of sedimentary amyloplasts/statoliths in columella cells decreased, and gravitropism of primary roots was partially reduced. Gene expression analyses showed that the expression of auxin response and biosynthesis genes *IAA1, IAA2, TAA1* and *YUC8* were repressed and the *SOS1* gene was upregulated two-fold in roots grown under moderate salt conditions. Proteomic analysis showed that co-chaperone and activator of HSPs such as Hop2 and Aha1 domain-containing protein orthologs were upregulated. Moreover, several secondary metabolic process-related proteins, antioxidant proteins, stress response proteins and proline catabolic process-related proteins were also increased. In contrast, enzymes associated with root hair elongation and nucleotide and protein syntheses were downregulated. These changes in auxin-related physiological responses, root architecture, lower ROS signaling, and stress-related protein expression promote primary root penetration into lower-salinity deeper soils as an adaptation of *S. parvula*.

## Introduction

Some plants can survive and grow in highly saline soil or water in natural ecosystems. Such plants, which can generally complete their life cycle with at least 200 mM NaCl, are classified as halophytes and constitute the natural vegetation of these areas (Rozema and Flowers, 2008, Flowers and Colmer 2008). During evolution, halophytes have developed various anatomical, physiological, biochemical, and molecular adaptations to withstand the detrimental effects of soil salinity. For this reason, halophytes offer an opportunity to elucidate new adaptation mechanisms and survival strategies that can be used to increase crop yields on saline soils.

Most of our knowledge of salt tolerance mechanisms comes from the model plant *Arabidopsis thaliana* (Orsini et al., 2010; Ozgur et al., 2013), which has not evolved to tolerate high soil salinity. So the search for new model plants that can tolerate high salinity was initiated in the family Brassicaceae to be able to exploit genetic tools available in Arabidopsis. *Schrenkiella parvula* (syn. *Thellungiella parvula* or *Eutrema parvulum*) is a halophyte that belongs to the same family as Arabidopsis and grows around Lake Tuz (Salt Lake) in central Turkey (Jarvis et al., 2014), one of the world’s largest hypersaline lakes (Oh et al., 2014). *S. parvula* was also named Arabidopsis-related model species (ARMS) because it can tolerate salt concentration of 600 mM NaCl (Orsini et al., 2010) and has 90% genome similarity to *A. thaliana* (Yalcinkaya et al., 2019). The genome and transcriptome analyses reveal a unique evolution of the Na^+^ extrusion and K^+^ uptake systems in reducing Na^+^ toxicity in *S. parvula*. For example, transcriptomic analysis demonstrated that *S. parvula* has higher basal and salt induced *SOS1* (Salt Overly Sensitive 1) expression when compared to Arabidopsis. Moreover, *S. parvula SOS1* promoter region was longer, and transcripts had a conserved pyrimidine-rich 5′ untranslated region. In addition, it has been demonstrated that HKT1;2 from *S. parvula*, which contains an amino acid substitution, can confer salt tolerance in Arabidopsis (Ali et al., 2018). Other studies that used *S. parvula* focused on wax production (Teusink et al., 2002), antioxidant defense (Uzilday et al., 2015), reactive carbonyl species metabolism (Yalcinkaya et al., 2019) and aldehyde dehydrogenases (Hou and Bartels, 2015) under salinity. In general, high concentrations of NaCl (>200mM NaCl) is used during these experiments to fully induce salt stress responsive mechanisms. However, little is understood about the physiological changes at moderate Na^+^ concentrations. This is an important consideration since the salinity of the rhizosphere in nature differs between the surface and the soil profile and can change greatly due to the moisture content of the soil (Hassanuzzaman et al., 2014). Therefore, plants have to adapt to low or high salinity along their growth trajectory.

In this work, we aimed to elucidate root growth strategies utilized by *S. parvula* at moderate salinity. To this end, the experiments were performed starting from moderate salt concentrations of 100 mM NaCl, showing that primary root elongation was inhibited at 200 mM, but conversely enhanced at 100 mM. Root growth analysis under mannitol-induced osmotic stress revealed that this response is specific to NaCl. Therefore, to further understand this strategy, we characterized physiological responses and determined changes in root anatomy, root cell division and expression of genes related to root growth at 100 mM NaCl. Finally, a proteomic approach was used to further elucidate response of root tissue to 100 mM NaCl at molecular level.

## Materials and Methods

### Plant growth conditions

Seeds of *S. parvula* and *A. thaliana* Col-0 were surface-sterilized with 0.5% Tween-20 and 5% NaClO for 10 min, washed three times with sterilized distilled water, and plated on 1/2 MS medium (1/2 MS Salt, 1% Sucrose, 0.7% Agar) (Murashige & Skoog, 1962). As part of the stratification process, plates were stored in the dark at 4 °C for 7 days. For germination and subsequent growth, plates were transferred to a 16 h light/8 h dark cycle at 23 °C in a plant growth chamber (MLR-352, PHCbi). Four-day-old seedlings were transferred to 1/2 MS medium containing different concentrations of NaCl or mannitol as indicated and subsequently grown in the same growth chamber.

### Observation of root architecture and cells

The whole image of the seedling root was taken with a digital camera (EOS Kiss X9, Cannon), and more detailed observation was performed with a stereomicroscope (SMZ25, Nikon) with a digital camera (DP73, Olympus). Epidermal cells in the root elongation zone were observed with a DIC microscope (BX51, Olympus), and internal meristems were observed with a confocal microscope (LSM 800, ZEISS) after 5□μM propidium iodide (PI, SigmaAldrich) staining. Amyloplasts of columella cells were observed with an optical microscope (BX51, Olympus) after staining with Lugol’s solution (SigmaAldrich). The Click-iT EdU Cell Proliferation Kit (ThermoFisher) was used for DNA replication activity in root cells, and 20□μM CM-H2DCFA (ThermoFisher) was used for monitoring intracellular reactive oxygen species (ROS). Each fluorescence image was taken with a confocal laser microscope (FV10i, Olympus and LSM 800, ZEISS) under the same excitation and exposure conditions for each sample. Primary root length, cell size, and fluorescent intensities were measured with Image J software (https://imagej.nih.gov/ij/).

### Real-time RT PCR analysis

Total RNA was isolated from dissected roots (around 10 samples) of 6-day-old seedlings cultured for 2 days in the presence or absence of 100 mM NaCl using TRIzol Reagent (Invitrogen). The expression analysis of each target gene was performed quantitatively by real-time RT-PCR method after cDNA synthesis using PrimeScript™ RT reagent Kit with gDNA Eraser (TaKaRa) and TB Green® *Premix Ex Taq*™ II (Tli RNaseH Plus) (TaKaRa) with Thermal cycler (CFX96, BioRad). The following primer pairs were used for PCR amplification: Tp7g12650 (*IAA1* ortholog) (forward: 5□-TCT TGC GAG AGA AGC AAC AA -3□, reverse: 5□-GAG ATA TGG AGC TCC GTC CA -3□), Tp3g20710 (*IAA2* ortholog) (forward: 5□-GGA TTA CCC GGA ACA GAG CA-3□, reverse: 5□-TTC GTC ACG GTT TTC CTC GT-3□), Tp5g25710 (*TAA1* ortholog) (forward: 5’-TGT AAG AGC GAG TCG GAG AAC-3’, reverse: 5’-CCT GCT CTG CTC ATG ACC TT-3’) Tp2g13410 (*SOS1*) (forward: 5□-ATG ACT TTG GGC ATG TTT TAT GCT GCA-3□, reverse: 5□-CCT TCA GCA ATG ACA ACA CCA CTG AGG A-3□), Tp6g18810 (*PIN2* ortholog) (forward: 5’-ATG ATC ACC GGC AAA GAC AT-3’, reverse: 5’-GCG TAA GGA TCA TTG GAG GA-3’), Tp5g00540 (*TIR1* ortholog) (forward: 5’-GTG GTG CAG GAA AAA GGT GT-3’, reverse: 5’-CTC CTC AAG CCA GGT GTA GG-3’), Tp7g26730 (*YUC8* ortholog) (forward: 5’-GCT TTG AGT TGG CTT GTG CT-3’, reverse: 5’-TTT CGC GCT CTT TAG CTC CA-3’), Tp1g12470 (*DAO1* ortholog) (forward: 5’-GAC GGA CTT GCA ATG GAT TT-3’, reverse: 5’-AGT CCG CCA ACA TTT TCA TC-3’), Tp1g12460 (*DAO2* ortholog) (forward: 5’-AGG CCT CTG CTG AAC AAA GA-3’, reverse: 5’-TCG TCA TCA CCA TGA ACG AT-3’) and Tg6g37750 (*UBC22* ortholog) (forward: 5□-GCT CGC CAA GGA ACT AAA GA-3□, reverse: 5□-ATG TGG AAA ATC ATG CGA CA-3□). The real-time RT-PCR data were normalized to Tg6g37750 (*UBC22*), which encodes an ortholog of ubiquitin-conjugating enzyme 22, as a housekeeping gene. The analyses were performed in at least biological triplicate for each sample.

### Proteome analysis

Total protein was extracted from the roots (15-25 samples) of 9-day-old seedlings cultured for 5 days in the presence or absence of 100 mM NaCl with a homogenizer (Micro Smash MS-100R, TOMY) using lysis buffer (7 M urea, 2 M thiourea, 5% CHAPS, 50 mM DTT, 0.2% Bio-Lyte) and zirconia beads. Extracted proteins were further purified using the BioRad ReadyPrep 2-D Cleanup Kit (BioRad), approximately 60-70 μg of protein was loaded onto 7 cm pH 3-10 NL and pH 4-7 IPG strips (BioRad) and subjected to first-dimensional electrophoresis with increasing voltages from 50V to 2000V. IEF strips equilibrated with buffers containing SDS and iodoacetamide (BioRad) were subjected to 2nd-dimensional electrophoresis by 10% SDS-PAGE. Protein spots was stained with Coomassie Brilliant Blue (CBB) and calculated as a percentage volume (% Vol) by Image Master 2D Platinum 6.0 software. Significant differences (*p* < 0.05) in expression levels were determined by at least three independent electrophoresis in each sample.

Each protein spot was excised from 2D SDS gels, digested with trypsin solution (Promega), and peptide fingerprinting was performed with a MALDI-TOF MS mass spectrometer (AB SCIEX 5800, SCIEX) and the MS-Fit ProteinProspector software (http://prospector.ucsf.edu/). MS/MS analysis was also performed by AB SCIEX 5800 and ProteinPilot Software (SCIEX). *S. parvula* protein sequence database was obtained from *Thellungiella parvula* genomic data and gene models (http://thellungiella.org/blast/db/TpV84ORFs.protein). NCBI-BLAST was used to identify their orthologues in *Arabidopsis*.

### Statistical analysis

The one-way ANOVA with post hoc Tukey’s HSD and Dunn’s test was used for comparisons between groups as appropriate (R or Origin software). All data points including outliers were used for means and statistical significance. Student’s *t* test was used for comparisons between two groups. A *p* value of < 0.05 was considered significant. Different letters indicate significant differences between the groups.

## Results

### Changes in root architecture in response to moderate salt stress in *S. parvula* seedlings

*S. parvula* 4-day old seedlings (primary root length approximately 1 cm) germinated on NaCl-free 1/2 MS medium were transferred and grown on media containing different concentrations of NaCl for 5 days. Interestingly, primary root elongation was significantly inhibited by 200 mM NaCl but conversely accelerated by 100 mM NaCl (Fig. 1 A, B, C). In contrast, root fresh weight decreased in a salt-dependent manner (Fig. 1D). Growth for an additional 5 days in medium containing 100 mM NaCl also resulted in significant elongation of primary roots, although they were thinner and had shorter lateral root lengths compared to 0 mM NaCl condition. (Suppl. Fig. 1).

**Figure 1.**
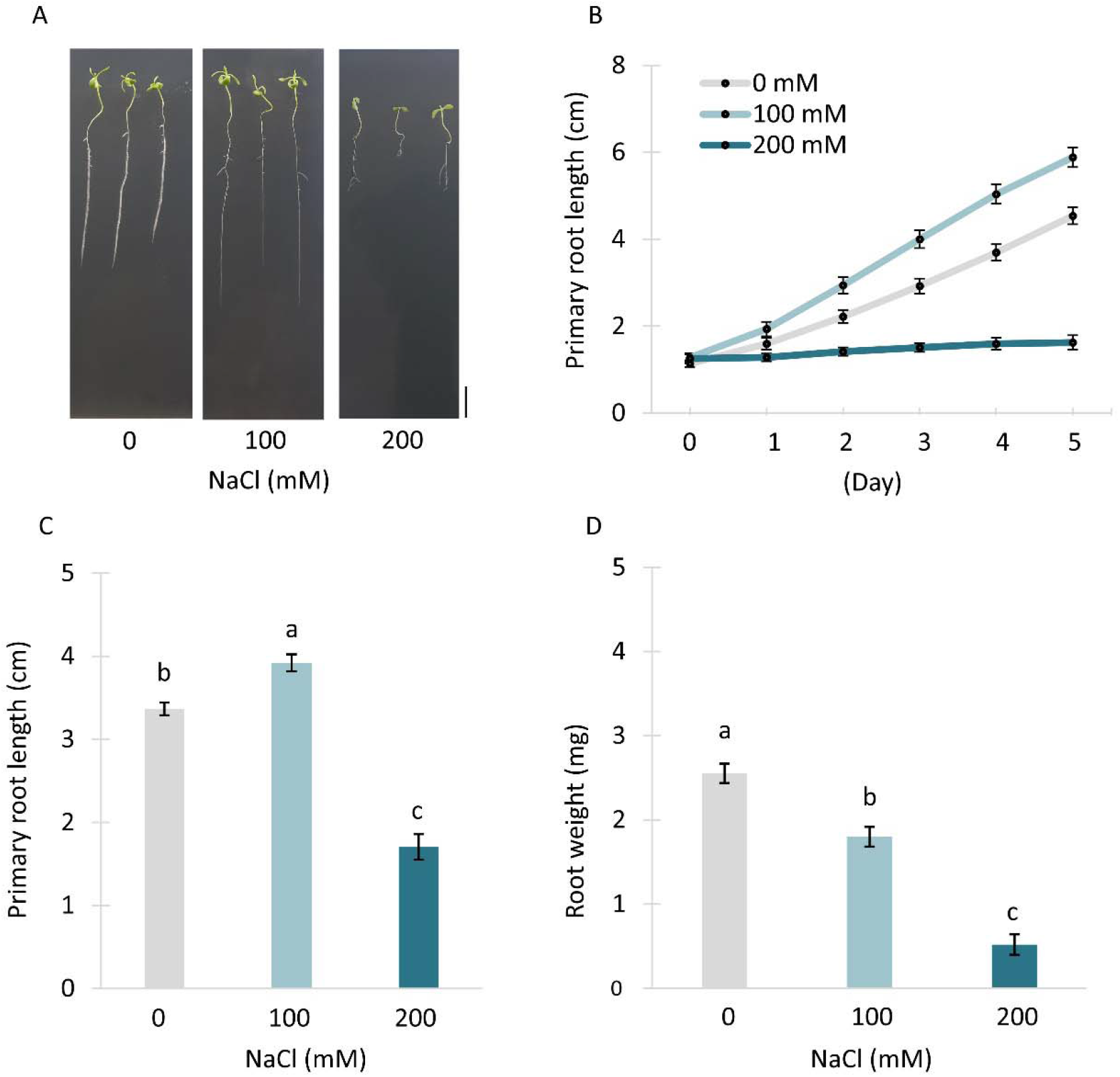
Root growth in *S. parvula* seedlings cultivated for 5 days on 1/2 MS medium containing different concentrations of NaCl. Four-day-old seedlings (post-germinated in NaCl-free medium) were transferred to 1/2 MS medium containing 0, 100, and 200 mM NaCl and cultured for an additional 5 days. (A) 9-day-old seedlings cultured for 5 days at the indicated NaCl concentrations. Scale bar: 1 cm. (B) Primary root growth under different NaCl conditions. (C) Primary root length in 9-day-old seedlings. (D) Fresh weight of primary root in 9-day-old seedlings. Groups were compared with one-way ANOVA Tukey post hoc test. Different letters indicate significant difference. Data represent *n* = 10 and means ± S.E.

For comparison with *A. thaliana*, seedlings of both species with a primary root length of about 0.5 cm germinated in NaCl-free medium were transferred to media with different salt concentrations and their growth was observed. Primary root growth of *Arabidopsis* seedlings was reduced by half under 100 mM NaCl conditions. Growth nearly stopped at 150 mM and died within 5 days above 200 mM NaCl (Suppl. Fig. 2). On the other hand, the primary roots of *S. parvula* seedlings elongated reproducibly even at the younger stage, and remarkably elongated under 100 mM NaCl condition. At 150 mM, the same level of elongation was observed as the 0 mM NaCl controls. Elongation was inhibited above 200 mM, but no death was observed at least below 400 mM NaCl (Suppl. Fig. 2).

**Figure 2.**
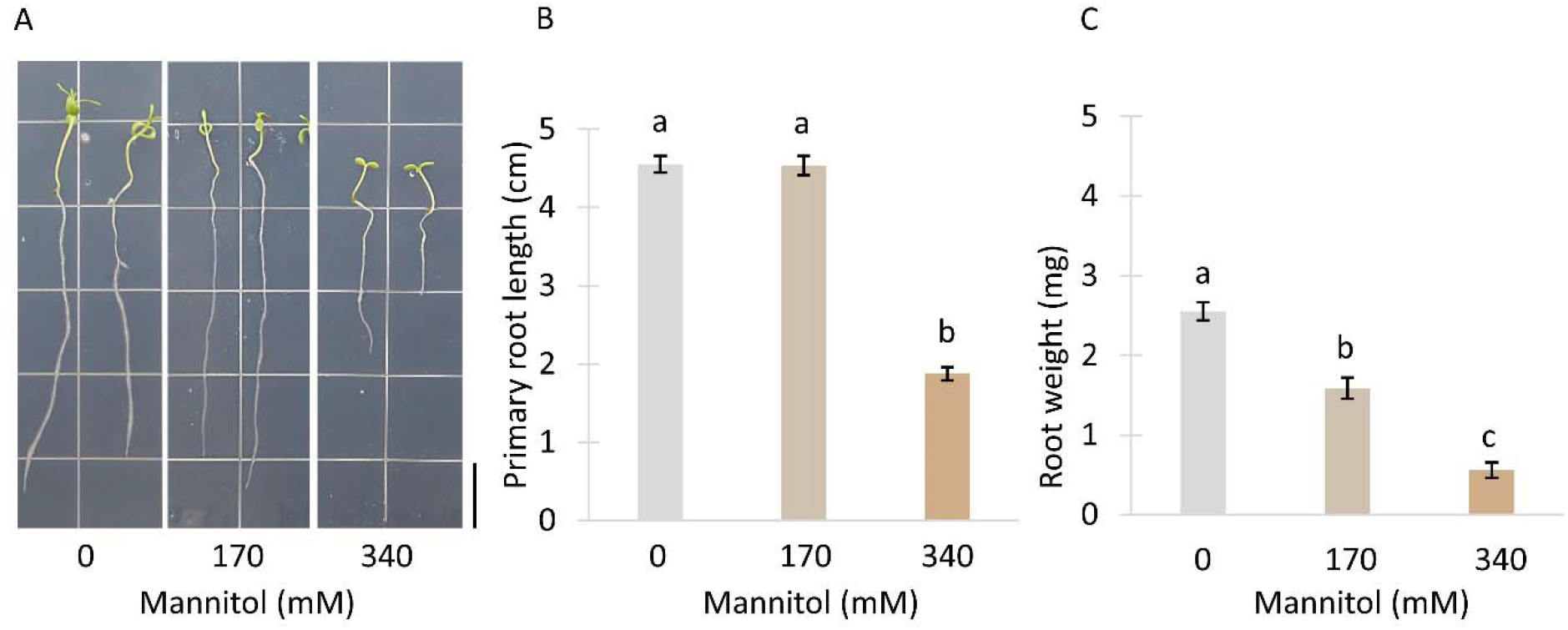
Root growth in *S. parvula* seedlings cultivated for 5 days on 1/2 MS medium containing different concentrations of mannitol. Four-day-old seedlings (post-germinated in NaCl-free medium) were transferred to 1/2 MS sucrose medium containing 0, 170, and 340 mM mannitol and cultured for an additional 5 days. (A) 9-day-old seedlings cultured for 5 days at the indicated mannitol concentrations. Scale bar: 1 cm. (B) Primary root length in 9-day-old seedlings. (C) Fresh weight of primary root in 9-day-old seedlings. Statistic test was used one-way ANOVA Tukey post hoc test. Different letters indicate significant difference. Data represent *n* = 10 and means ± S.E.

On the other hand, 170 mM mannitol, which is iso-osmotic with 100 mM NaCl (Cuin and Shabala, 2005), did not promote primary root elongation, but fresh weight decreased similarly to salt conditions. Further growth impairment was observed at 340 mM mannitol (Fig. 2). Furthermore, very interestingly, root hair formation was also most inhibited in the 100 mM NaCl condition compared to the 200 mM NaCl, 170 mM mannitol and 340 mM mannitol conditions (Fig. 3). These results clearly demonstrate that primary root elongation of *S. parvula* seedlings under moderate salt stress conditions of 100 mM NaCl is specific to ionic effects rather than osmotic stress avoidance responses. This might be a strategy of *S. parvula* roots to rapidly reach deeper soil with low salinity under moderate salt stress.

**Figure 3.**
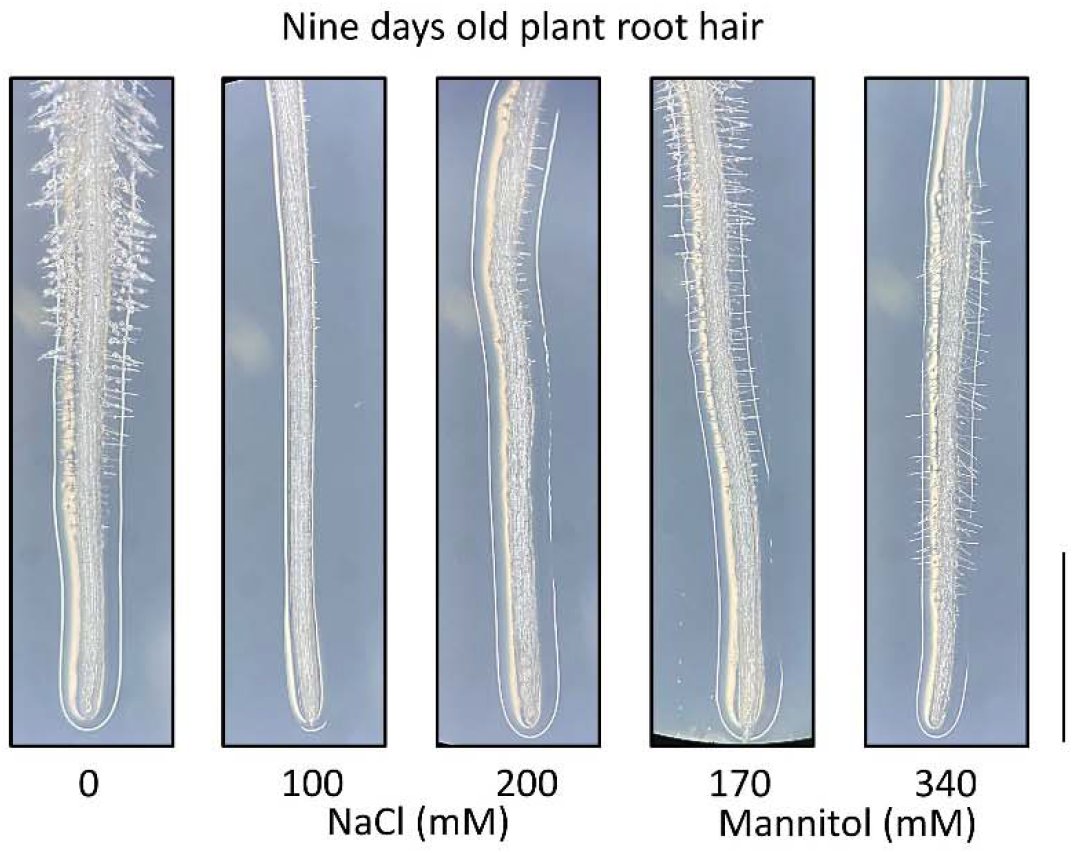
Root hair formation in 9-day-old *S. parvula* seedlings cultivated for 5 days on 1/2 MS medium containing different concentrations of NaCl and mannitol. Represents a typical image of a root apical part. Scale bar: 1 mm.

### Decreased meristematic DNA replication under moderate salt stress

To understand the mechanism by which primary root elongation of *S. parvula* seedlings is accelerated under moderate salinity conditions, we measured root meristem size and cell length in the transitional elongation zone after transfer to medium containing 100 mM NaCl. Analysis of root epidermal cells by differential interference contrast microscopy (DIC) revealed that those cells exposed to moderate salinity for 5 days elongated faster in the elongation zone (Fig. 4 A, B). The average length of the 8th epidermal cell after elongation initiation was approximately 100 μm in salt conditions, 1.4 times longer than in controls (70 μm). Root tip meristematic cells were stained with propidium iodide (PI) and observed with a confocal microscope. A significant reduction in meristematic tissue size was already observed on day 2 of moderate salt treatment (Fig. 4 C, D). Furthermore, 100 mM salt damaged some basal meristematic cells such that PI penetrated into the cells. On the other hand, in Arabidopsis seedlings, a further increase in these damaged dead cells was observed not only in basal meristematic cells but also in the apical meristematic cells (Suppl. Fig. 3).

**Figure 4.**
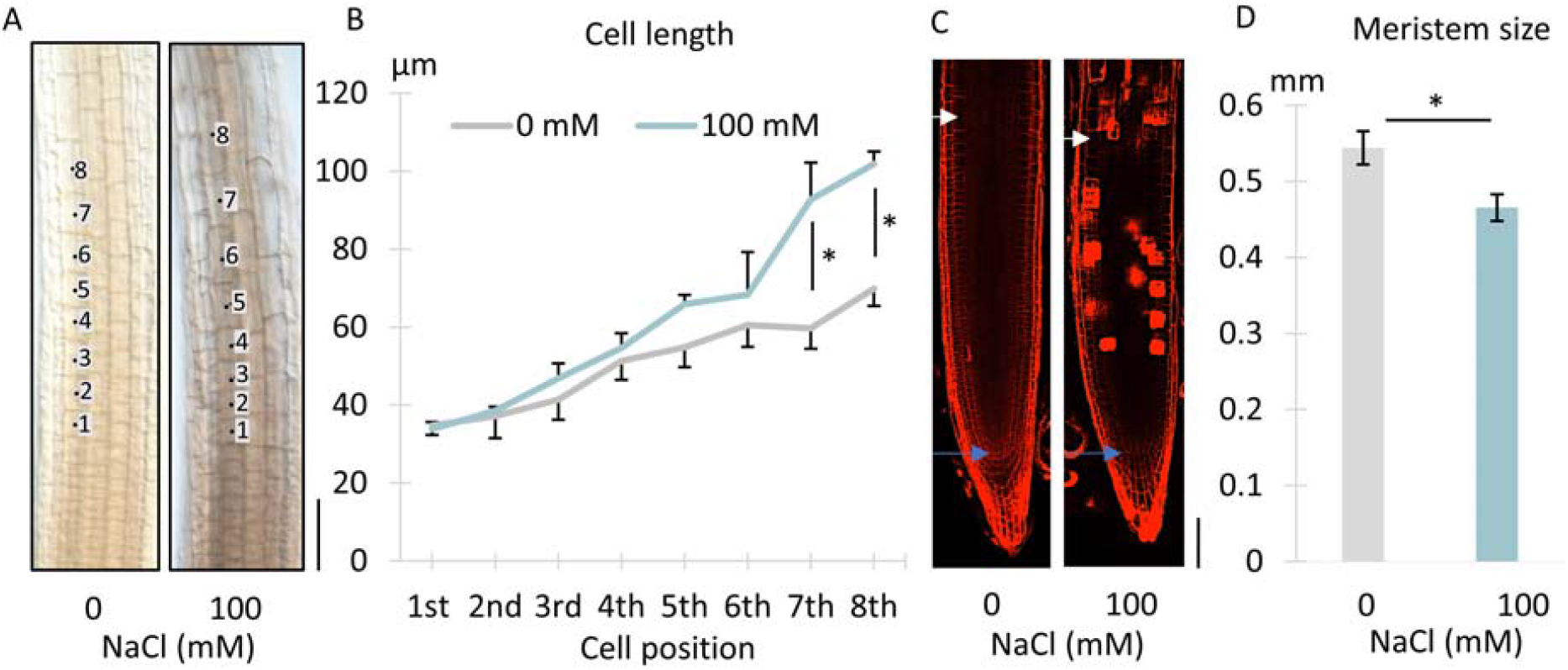
Epidermal cell elongation and meristem size in roots of *S. parvula* seedlings under moderate salinity. Four-day-old seedlings were transferred to 1/2 MS medium containing 0 and 100 mM NaCl and cultured for indicated period. (A) DIC image of root epidermal cells in elongation zone of 9-day-old seedlings cultured in 0 and 100 mM NaCl for 5 days. (B) Average length of epidermal cells from the beginning of elongation to the eighth. Data represents *n* = 3 and mean ± S.E. Student’s *t* test * *p* < 0.05. (C) Confocal microscopy image of PI-stained root meristems of 6-day-old seedlings cultured in 0 mM and 100 mM NaCl for 2 days. Some dead cells were visualized in the basal meristem area. Blue arrows: quiescent center. White arrows: elongated cells. Scale bar: 100 μm. (D) Average length of root apical meristem. Data represent *n* = 7 and mean ± S.E. Student’s *t* test * *p* < 0.05. Scale bars: 100 μm.

To further understand the effects of moderate salinity on cell proliferation in root meristems, we monitored DNA replication activity by incorporation of 5-ethynyl-2’-deoxyuridine (EdU). Four-day-old seedlings grown under NaCl-free conditions were further cultured in 100 mM NaCl for 2 days, followed by root infiltration with EdU for 4 h. In NaCl-free control root meristems, approximately 900 nuclei were visualized with EdU (Fig. 5A). On the other hand, the 100 mM NaCl condition reduced EdU-incorporated nuclei in the root meristem by 26% (Fig. 5B, C). Moreover, among three levels of fluorescence intensity for each nucleus, we observed that the ratios of strongest and intermediate nuclei were significantly reduced, leading to a large decrease in DNA replication activity (Fig. 5C). These results clearly indicated that moderate salinity-induced primary root elongation was due to enhanced cell elongation in the elongation zone rather than activation of root meristematic cell proliferation.

**Figure 5.**
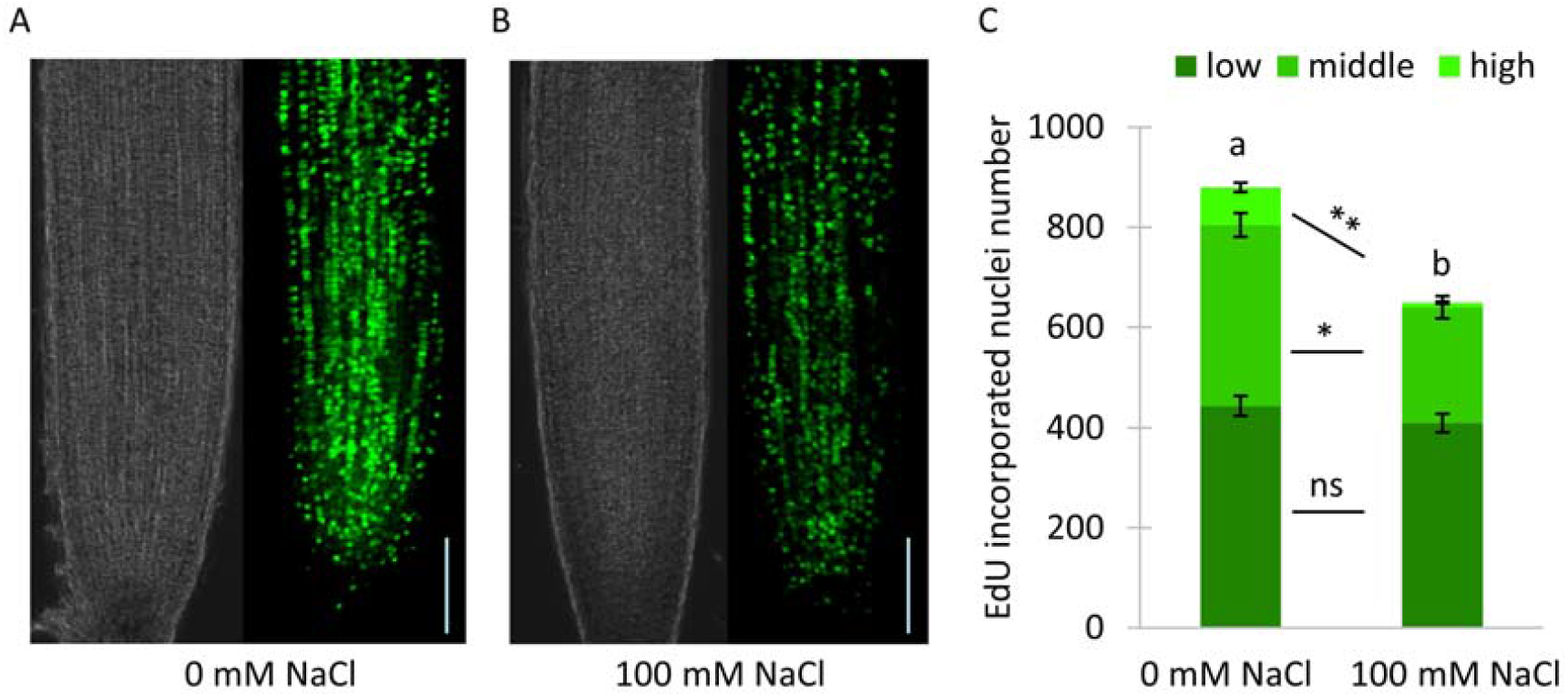
Incorporation of EdU in root meristem of 6-day-old *S. parvula* seedlings cultured in 0 mM and 100 mM NaCl for 2 days. (A) Phase-contrast (left) and EdU fluorescent (right) confocal microscopy images of root meristems cultured in 0 mM NaCl. (B) Same at 100 mM NaCl. Each fluorescence images were taken under the same excitation and exposure conditions for each sample. Scale bars: 100 μm. (C) Number of nuclei with EdU fluorescence classified at three different intensities. Data represent *n* = 7 and mean ± S.E. Statistic test was used Dunnett’s test for each category (* *p* < 0.05, ** *p* < 0.01, ns: not significant) and one-way ANOVA Tukey post hoc test for total nuclei number (different letters indicate significant difference).

### Decrease in auxin signaling and increase in ROS levels under moderate salinity

On the surface of 1/2 MS medium with moderate salt, *S. parvula* roots tended to bend randomly compared to controls (Suppl. Fig. 1). Therefore, we investigated the gravitropism of seedling roots grown under 100 mM NaCl conditions for 2 days, and observed the statoliths / amyloplasts of root columella cells. The results shown in Fig. 6 indicate that growth in the presence of moderate salt suppressed the size of amyloplasts and reduced the starch content to 40% compared to controls.

**Figure 6.**
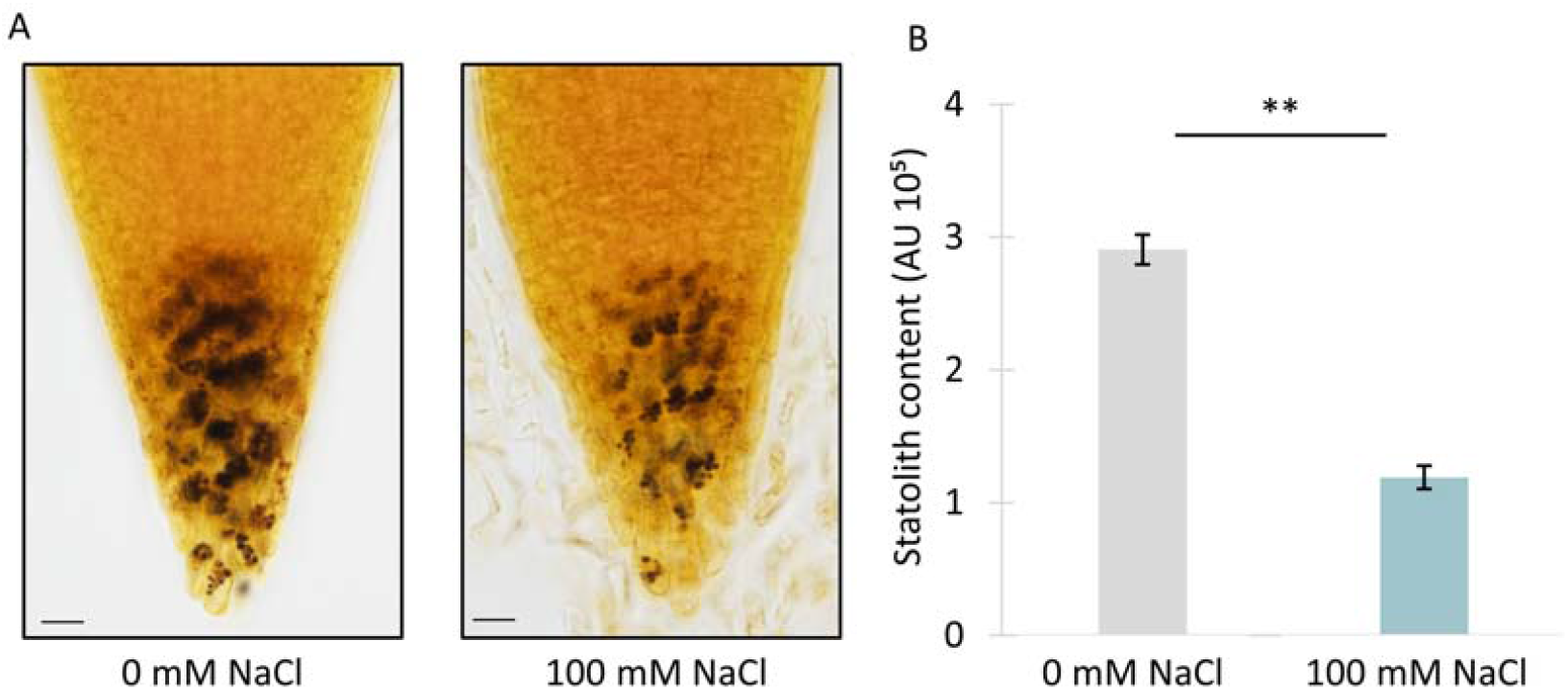
Amyloplasts of columella cells in 6-day-old *S. parvula* seedlings cultured in 0 and 100 mM NaCl for 2 days. (A) Amyloplasts were stained with Lugol’s solution and observed with an optical microscope. Scale bars: 20 μm. (B) Statolith amount was calculated by Image J software. Data represent *n* = 10 < and mean ± S.E. Student’s *t* test ** *p* < 0.01.

As a control, 4-day-old seedlings were transferred to fresh medium without NaCl and grown for 2 days before being tilted horizontally or held vertically. After 24 hours, root elongation was accelerated by 15% by tilting horizontally compared to remaining vertical, but not significantly (Table 1). The average flexion angle due to gravitropism was 66° ± 2.5° (Table 1). On the other hand, the seedlings were similarly transferred to a medium containing 100 mM NaCl, grown for 2 days, and tilted horizontally. After 24 h, root elongation did not change compared to seedlings that remained vertical. The bending angle due to gravitropism was 57° ± 2.2°, indicating a small but significant decrease in root gravitropism at moderate salinity (Table 1, Suppl. Fig. 4).

**Table 1.** Twenty-four-hour gravity response of roots in 6-day-old *S. parvula* seedlings cultured for 2 days under moderate salinity

|  | Position | Elongation length<br>(mm for 24 hours) | Bending angle<br>(degree ° for 24 hours) |
| --- | --- | --- | --- |
| 0 mM NaCl | Vertical | $5.8 \pm 0.05^b$ | $4.3 \pm 1.7^a$ |
| | Horizontal | $6.6 \pm 0.07^b$ | $66 \pm 2.5^b$ |
| 100 mM NaCl | Vertical | $8.2 \pm 0.08^a$ | $2.9 \pm 1.2^a$ |
| | Horizontal | $8.8 \pm 0.07^a$ | $57 \pm 2.2^c$ |

Statistic test was used one-way ANOVA Tukey post hoc test for elongation length and bending angle, respectively. Different letters indicate significant difference.

Not only primary root elongation, but also root hair formation, gravitropism, and statoliths/amyloplasts are controlled by auxin signaling (Baldwin et al. 2013; Zhang et al. 2019). Therefore, the effects of moderate NaCl on the expression of genes related to auxin response, biosynthesis and transport were examined at steady state after 2 days of treatment. Comparing the relative expression levels with Tp6g37750 (*UBC22*) gene, the expression of Tp2g13410 (*SOS1*) gene increased approximately two-fold. Indeed, the expression of auxin response and biosynthesis genes, Tp7g12650 (*IAA1*), Tp3g20710 (*IAA2*), Tp5g25710 (*TAA1*), and Tp7g26730 (*YUC8*), was downregulated by approximately half in seedling roots grown in 100 mM NaCl (Fig. 7). In contrast, the expression of auxin transport related genes, Tp6g18810 (*PIN2*) and Tp5g00540 (*TIR1*), was slightly increased (Fig. 7). The expression levels of the auxin catabolic genes Tp1g12470 (*DAO1*) and Tp1g12460 (*DAO2*) were nearly consistent under moderate salinity. These results strongly suggest that moderate salt stress may reduce auxin levels in *S. parvula* roots through suppression of auxin biosynthetic pathways. Therefore, we exogenously added auxin NAA (final concentration 1 μM) to seedlings grown in the presence of NaCl and observed the root morphology. As a result, NAA significantly inhibited primary root elongation and restored root hair formation (Fig. 8). However, in the presence of 100 mM NaCl, primary roots were still longer and had fewer root hairs than in the case of adding NAA to the control.

**Figure 7.**
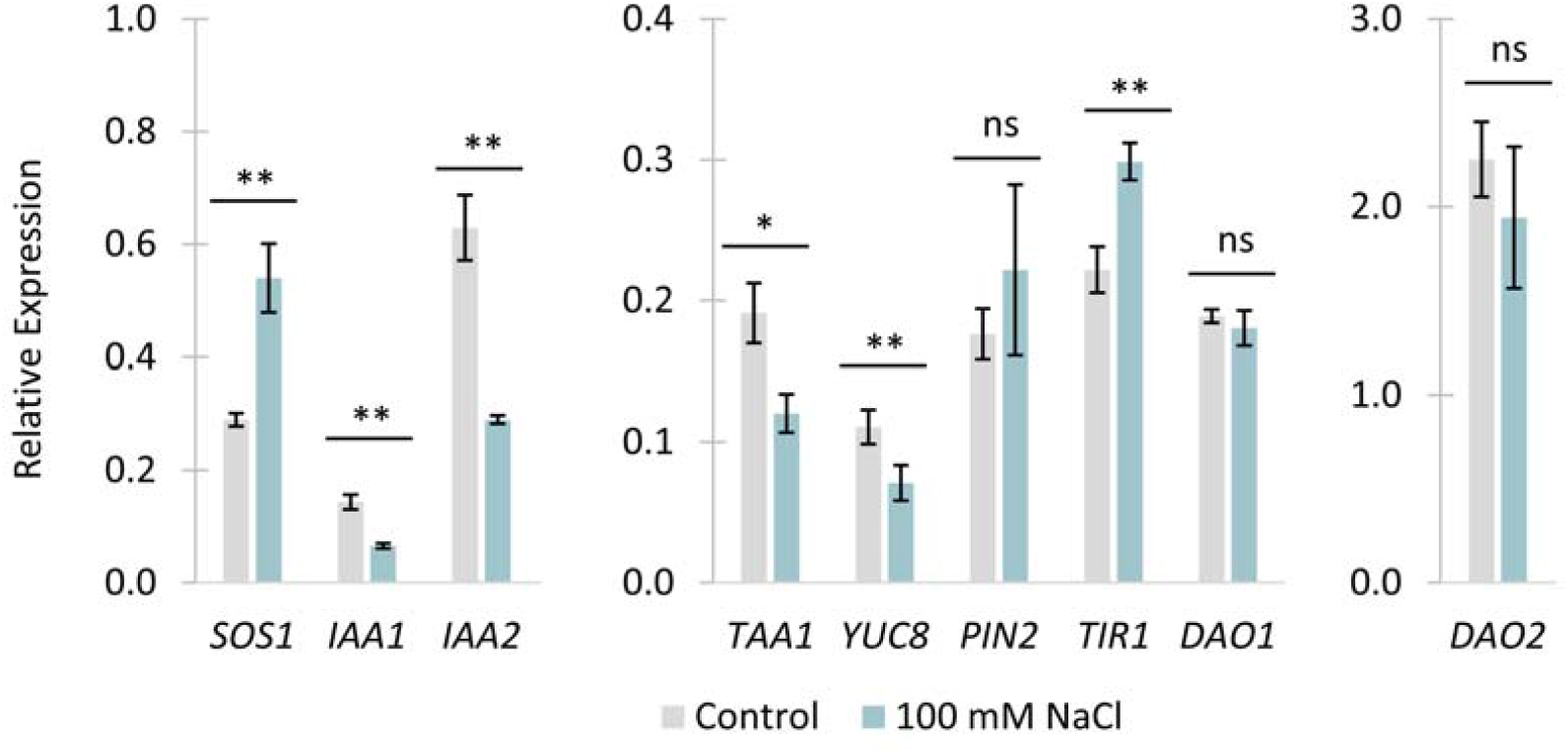
Relative expression level of auxin related genes (*IAA1, IAA2, TAA1, YUC8, PIN2, TIR1, DAO1*, and *DAO2*) and *SOS1* in roots of 6-day-old *S. parvula* seedlings cultured in 0 mM and 100 mM NaCl for 2 days. Real-time quantitative RT-PCR analysis was performed using gene-specific primers and relative expression ratios were calculated using *UBC22* gene expression. Data represent at least three biological samples and mean ± S.E. Student’s *t* test * *p* < 0.05, ** *p* < 0.01, ns: not significant).

**Figure 8.**
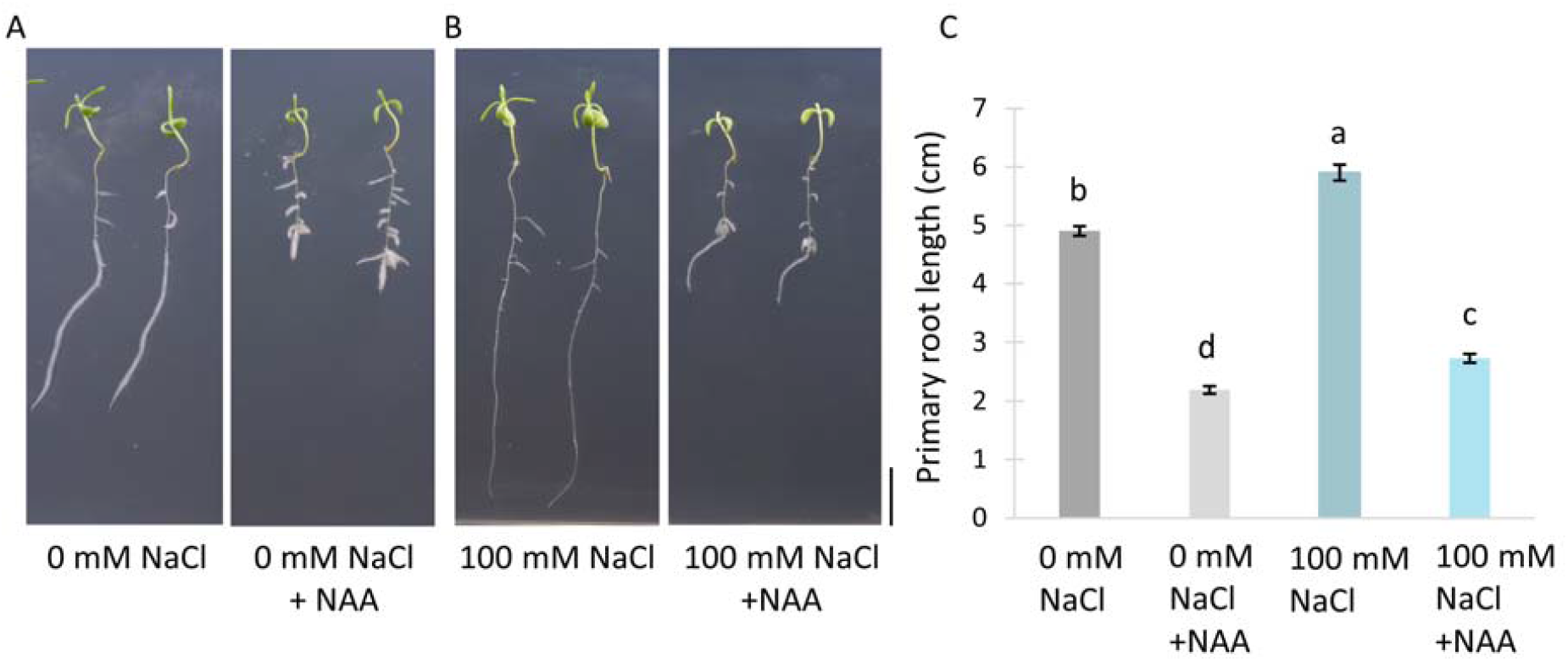
Inhibition of *S. parvula* primary root elongation by 1 μM NAA application. Four-day-old seedlings were transferred to 1/2 MS medium containing 0 and 100 mM NaCl in the presence or absence of 1 μM NAA and cultured for an additional 2 days. (A) Typical images of 6-day-old seedlings cultured in 0 mM NaCl. (B) Same at 100 mM NaCl. (C) Primary root length in 6-day-old seedlings. Groups were compared witg one-way ANOVA Tukey post hoc test. Different letters indicate significant difference. Data represent *n* = 6 and means ± S.E.

In addition to hormonal regulation, biotic and abiotic stress conditions produce reactive oxygen species (ROS) in plants (<u>Mittler et al., 2022</u>). We therefore monitored the ROS levels in roots of *A. thaliana* and *S. parvula* grown under 100 mM NaCl conditions. In controls, fluorescent signals using CM-H2DCFDA, a general oxidative stress indicator, was higher in *S. parvula* basal region compared to *A. thaliana*, and it increased significantly with differentiation beyond the elongation zone (Fig. 9). Under 100 mM NaCl conditions, there was a 4-fold increase in fluorescence signal in the apical meristem region of *S. parvula*, but no change in the basal region. In Arabidopsis, on the other hand, we observed a significant increase in fluorescence signal of >10-fold in both the apical and the basal region beyond the elongation zone. These results clearly show that 100 mM NaCl-induced ROS were much higher in *A. thaliana* than in *S. parvula*.

**Figure 9.**
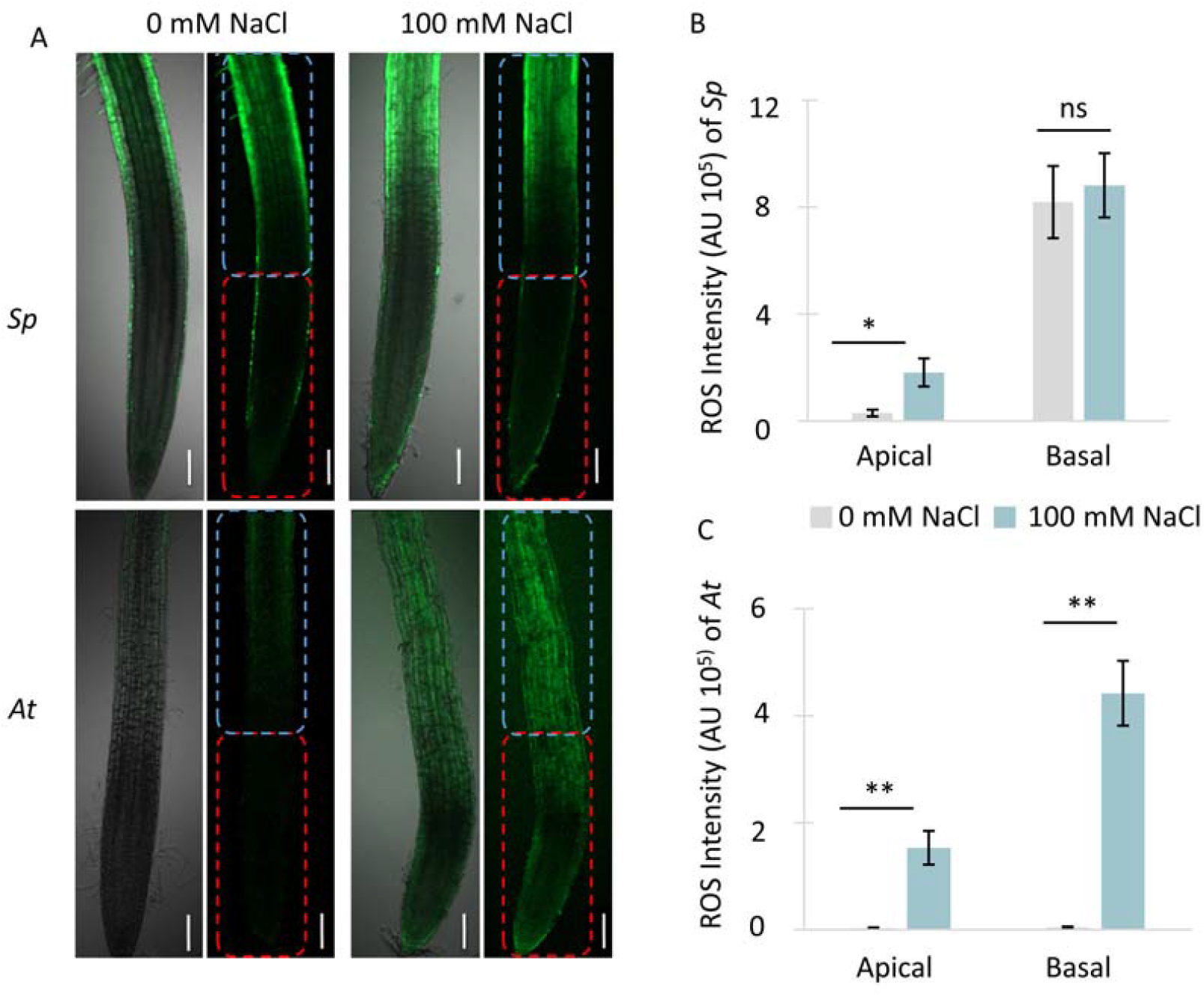
ROS induction in roots of 6-day-old *S. parvula* and *A. thaliana* seedlings cultured in 0 and 100 mM NaCl for 2 days. (A) Seedling roots of *S. parvula* (upper) and *A. thaliana* (bottom) were observed with CM-H2DCFDA fluorescence and phase contrast (left panels) and fluorescence only (right panels). Fluorescence signal intensity was measured for each 600 μm region (Apical: red square, Basal: blue square) in (B: *S. parvula*) and (C: *A. thaliana*). Each fluorescence images were taken under the same excitation and exposure conditions for each sample. Scale bars: 100 μm. Data represent *n* = 7 and mean ± S.E. Student’s *t* test * *p* < 0.05, ** *p* < 0.01, ns: not significant.

### Proteomic analysis of moderate salt stress-responsive proteins

To better understand the molecular response of 9-day-old seedling roots to moderate salinity (see Fig. 1A), extracted proteins were separated by 2D gel electrophoresis (2DE) and identified by MALDI-TOF analysis. Total proteins were separated at least three times using two isoelectric focusing methods (pI 4-7 and pI 3-10), respectively. A total of 698 protein spots were reproducibly resolved in the control and 605 protein spots in 100 mM NaCl (Suppl. Fig. 5). By MALDI-TOF analysis, we identified the presence of low-molecular-weight spots due to degradation and spots with different pI due to some modification, and finally identified a total of 300 different proteins (Suppl. Tables 1 and 2).

Of these identified proteins, 45 proteins were significantly changed in response to moderate salinity (Fig. 10, Table 2). Among them, 25 proteins showed a significant decrease and 20 proteins showed a significant increase. Furthermore, Figure 11 showed that among upregulated proteins GO enrichment analysis using the Arabidopsis orthologue revealed significant GO term enrichment for proteins involved in secondary metabolic process: Tp3g27850 (PRX32), Tp7g31780 (CCoAOMT1), Tp4g29660 (GSTF8), Tp4g13320 (GSTF9), and Tp5g33470 (GSTU19); response to oxidative stress: Tp3g27850 (PRX32), Tp7g32860 (CAT2), Tp5g28230 (KTI1) and Tp5g33470 (GSTU19); salt stress: Tp3g27850 (PRX32), Tp2g25700 (P5CDH), and DN9862 (PYK10); protein folding: Tp2g01900 (Hop2) and Tp3g10280 (Aha1-domain containing protein) (Table 2). In contrast, downregulated proteins were involved in purine nucleotide biosynthetic process: Tp7g33150 (AT4G35360), Tp6g12990 (MAB1), Tp1g23690 (APT1), and Tp6g39440 (ADK2); root hair elongation: Tp2g08860 (AT5G44020) and Tp6g33600 (ACT7); and translation elongation: Tp2g05140 (AT1G07920) and Tp6g24890 (AT5G19510) (Table 2). Interestingly, the proline catabolic enzyme, delta-1-pyrroline-5-carboxylate dehydrogenase Tp2g25700 (P5CDH) was upregulated, and conversely, glutamate dehydrogenase Tp6g26150 (GDH1) and glutamine synthetase Tp7g06630 (GLN2) were downregulated, suggesting that proline content would decrease and glutamate content would increase.

**Table 2.**
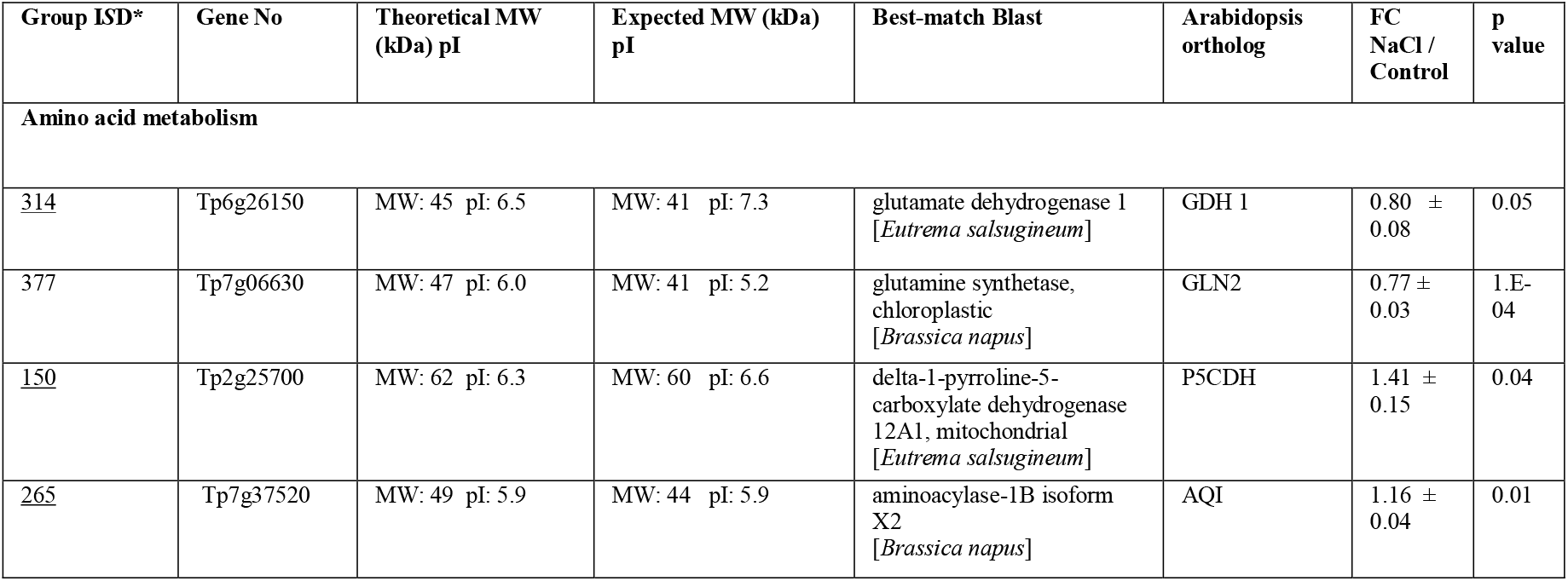
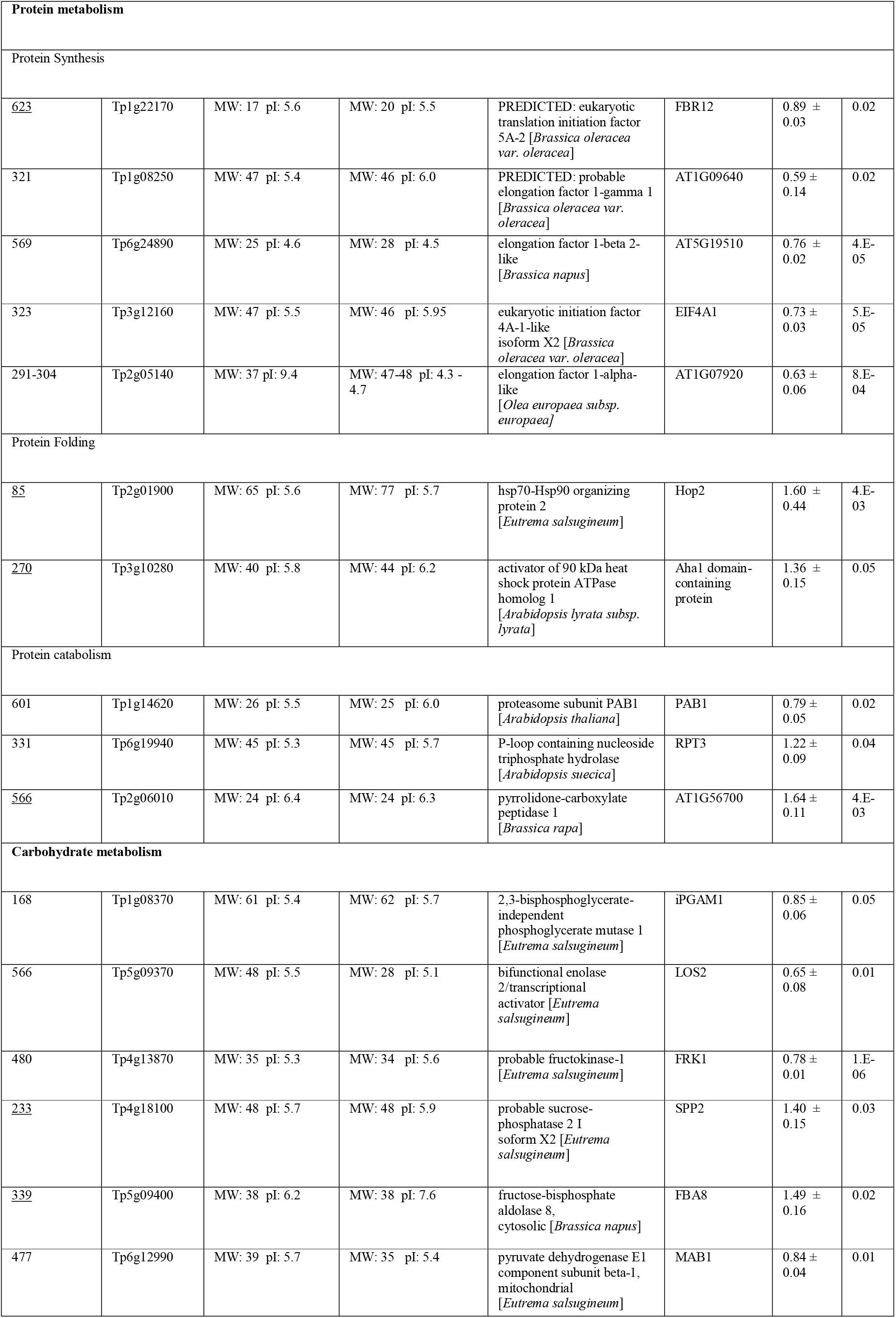
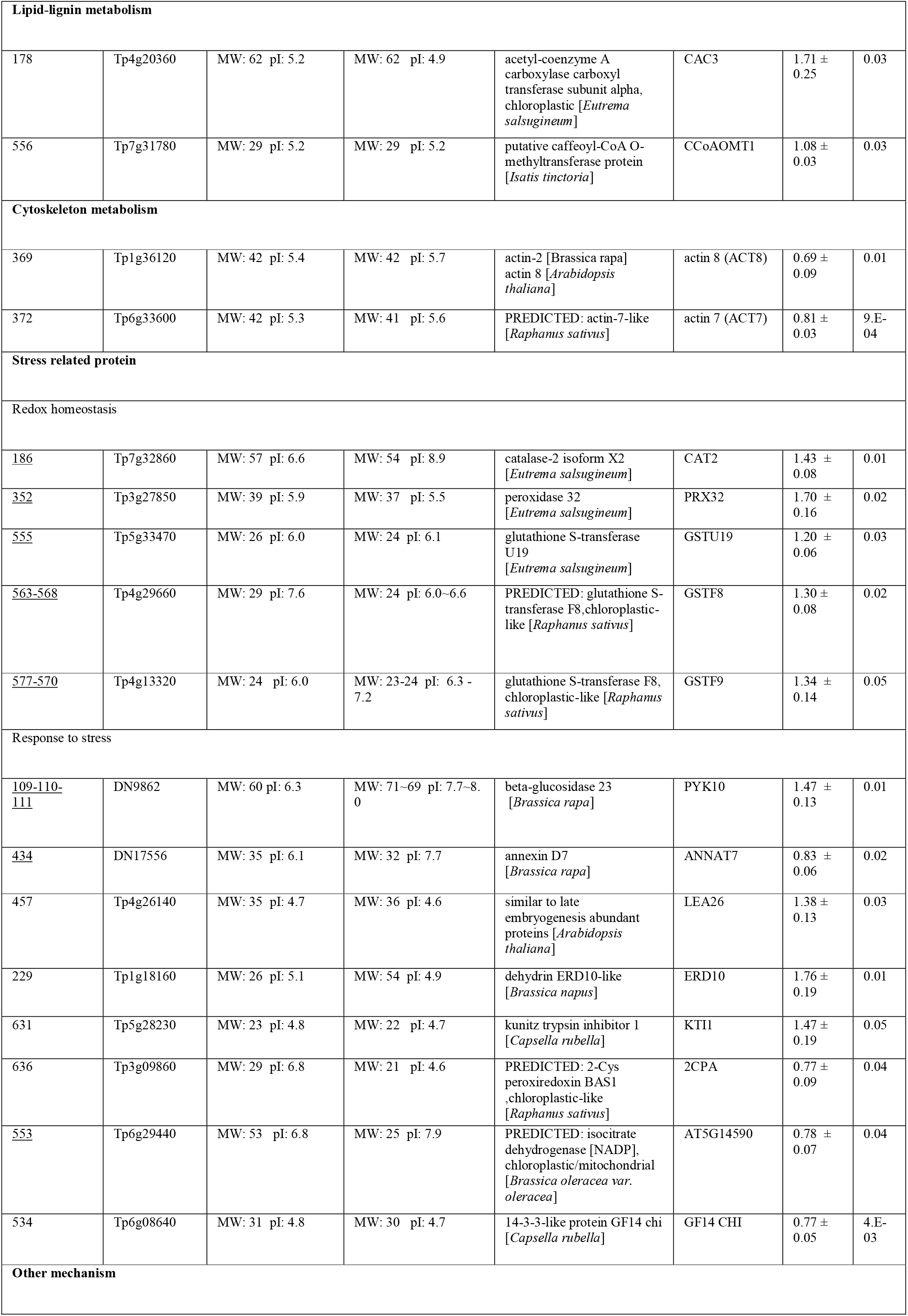
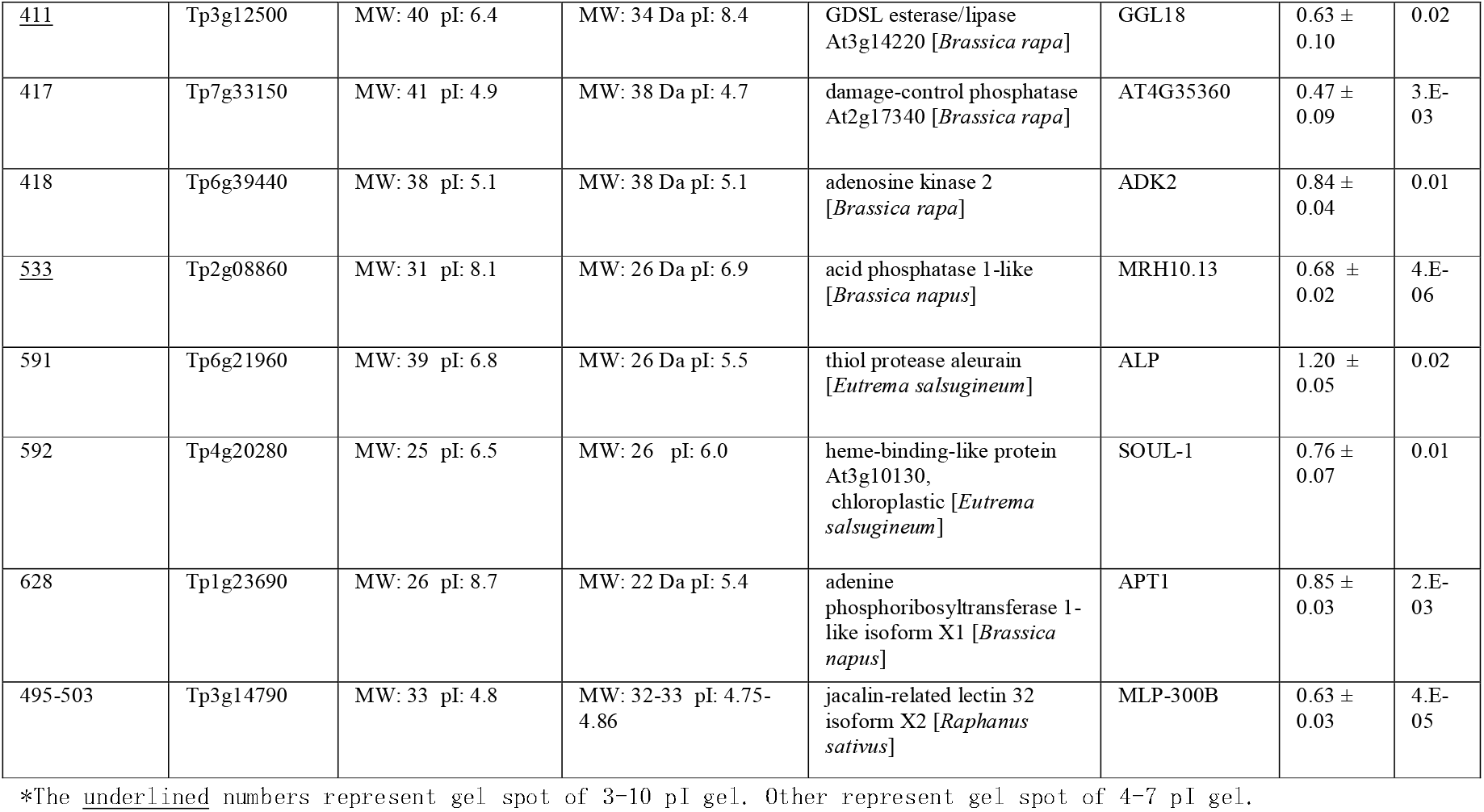
Identification of differentially expressed proteins in roots of *S. parvula* seedlings grown under 100 mM NaCl conditions from 2D-gel analysis

**Figure 10.**
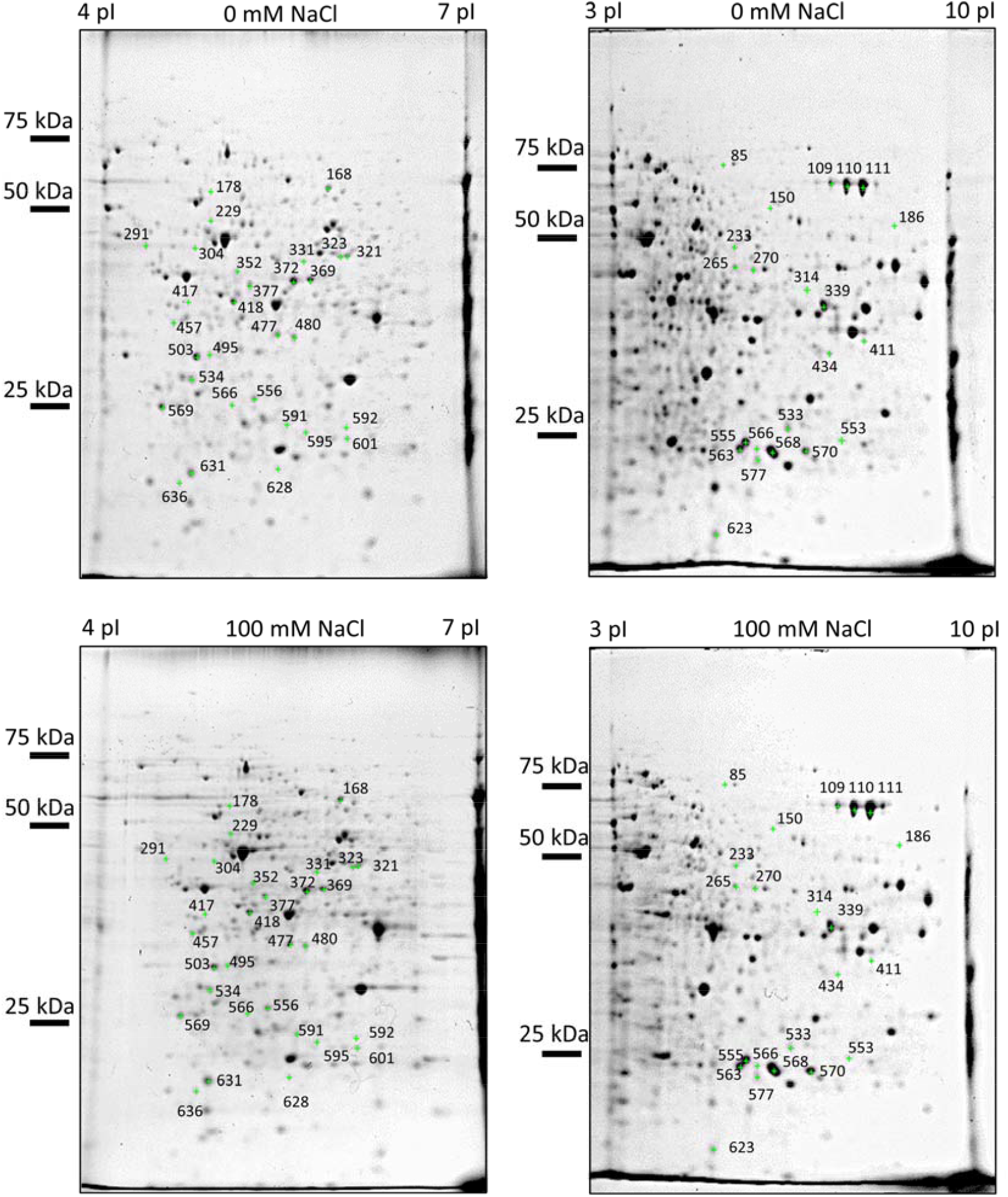
2DE images of total protein extracted from roots of 9-day-old *S. parvula* seedlings cultured in 0 mM and 100 mM NaCl for 5 days. pH 3-10 NL (left panels) and pH 4-7 (right panels) IPG strips were used for first dimensional electrophoresis. Numbers indicate group IDs of differentially expressed proteins shown in Table 2.

**Figure 11.**
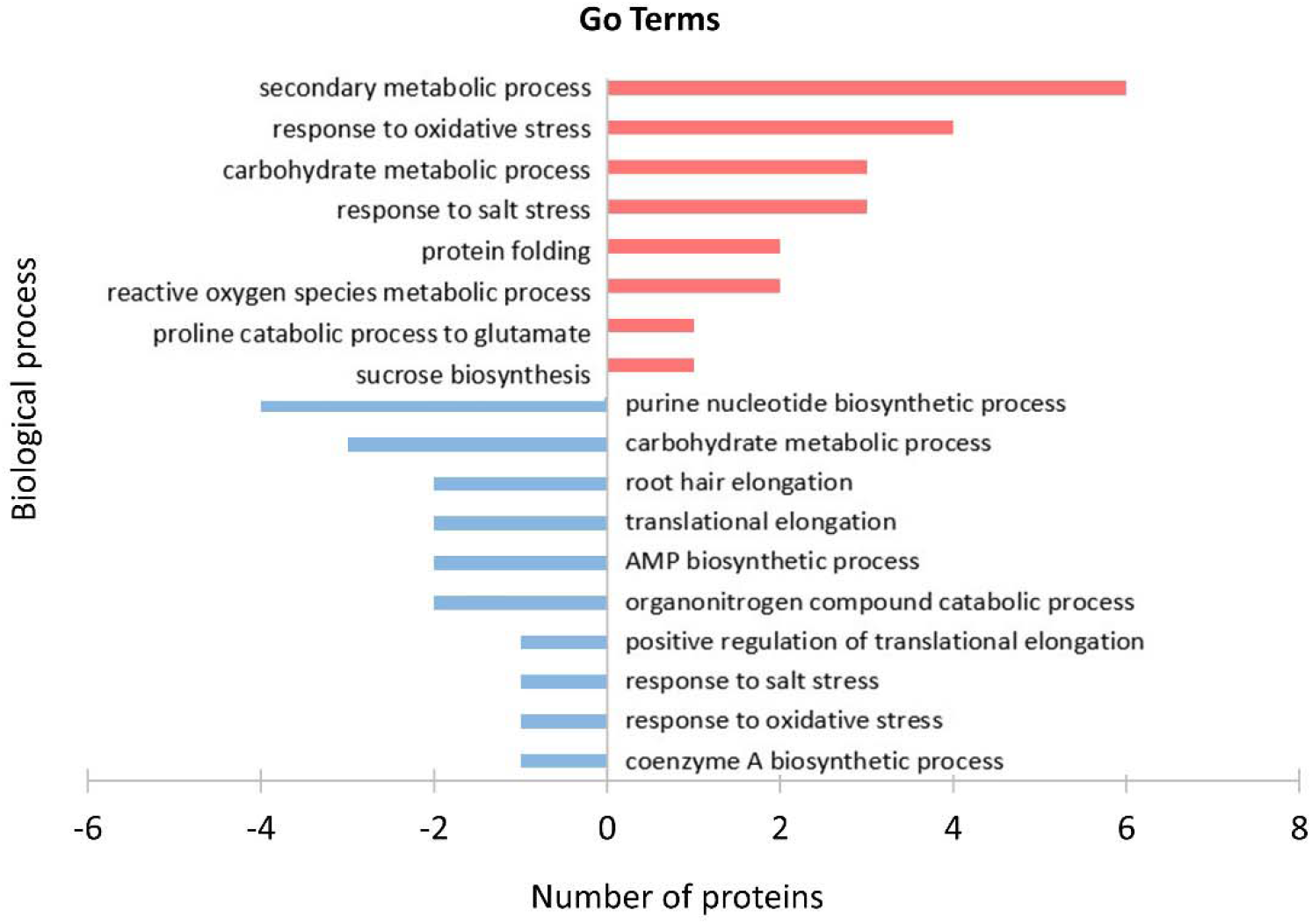
GO enrichment analysis of differentially expressed proteins in roots of 9-day-old *S. parvula* seedlings cultured in 0 mM and 100 mM NaCl for 5 days. Bar plot showing GO enrichment analysis (Panther, FDR < 0.05) for biological process in 45 differentially expressed proteins shown in Table 2.

## Discussion

Plant root architecture exhibits exceptional plasticity in response to changes in soil conditions and this regulation is a very important aspect of plant stress adaptation (Dinneny 2019). Plant root responses to soil water potential (hydrotropism and hydropatterning) and soil salinity (halotropism) and nutrient availability are well-documented phenomena (Hodge et al., 2009). In Arabidopsis, it has been demonstrated that mild NaCl concentrations (up to 50 mM NaCl) can increase the proliferation of lateral roots with a reduction in total lateral root and primary root length (Zolla et al., 2010). On the other hand, higher concentrations of NaCl leads to decreases in number of lateral roots. In addition, examination of root phenotypes of four different Arabidopsis ecotypes showed that NaCl concentrations above 50 mM NaCl results in serious inhibition of root growth in terms of primary root and lateral root length and lateral root number (Julkowska et al., 2014). However, induction of primary root length in Arabidopsis during salinity is not a documented phenomenon. Results of our study indicates that *S. parvula* utilizes a completely different strategy to regulate its root growth under mild salinity when compared to Arabidopsis, in which *S. parvula* favors primary root elongation under mild salinity (100 mM NaCl). This might have various implications in its natural habitat. *S. parvula* resides in a closed basin area that has high surface evaporation (Orhan et al., 2019). Therefore, movement of ions to surface due to capillary action is among major factors of surface salt accumulation and deposition (Rose et al., 2005). In such soils, Na levels tends to decrease with soil depth. Therefore, it might be beneficial for *S. parvula* to elongate its primary root to reach soil profile with lower salinity. After observation of increased primary root length upon 100 mM NaCl treatment, we aimed to elucidate the underlying mechanisms. Interestingly, root elongation was not observed at iso-osmotic mannitol concentrations, indicating that primary root elongation is a response to NaCl ions effects rather than to a decrease in water potential or an increase in osmotic stress.

General root elongation relies on a proper balance between cell proliferation and differentiation. In the root tip tissue, meristem growth is the result of cell division in the proximal meristem and cell differentiation in the elongation and differentiation zones (Sozzani and Lyer-Pascuzzi, 2014). Especially, root meristem size is controlled by hormonal regulation and ROS (Sozzani and Lyer-Pascuzzi, 2014). Among them, auxin, which crosstalks with cytokinin, interacts antagonistically at root transition zone (TZ) to balance cell differentiation with cell division to control root growth and elongation (Kong et al., 2018). In addition, auxin increase length of root hairs and number of lateral roots in a dose-dependent manner (Ishida et al., 2008; Peret et al., 2009). Moreover, we observed, reduced meristematic cell number, decreased DNA replication activity, and suppression of lateral root and root hair formation in *S. parvula* seedlings at moderate salt concentrations in addition to decreased expression of auxin-inducible genes. Proteomic analysis showed that moderate salinity upregulated class III peroxidase 32 (PRX32) might be involved in cell wall biosynthesis and auxin catabolism in *S. parvula* seedling roots. Recently, Liu et al. (2021) also reported significant increases in PRX32 and PRX34 in *Arabidopsis* salt stress-sensitive mutants (Liu et al., 2021). All these results strongly suggested that the root auxin signal in *S. parvula* was reduced under moderate salinity conditions.

ROS balance of hydrogen peroxide (H_2_O_2_) and superoxide (O_2_·−) between cell proliferation and cell elongation zones also regulates meristem size in an auxin-independent manner in Arabidopsis seedling roots (Tsukagoshi et al., 2010). Increased H_2_O_2_ levels in the apical region compared to O_2_·− lead to suppression of meristem size. In this study, we found that 100 mM NaCl treatment in *S. parvula* roots slightly but significantly increased ROS in the apical meristem region but not in the elongation and differentiation zones. Future studies should clarify whether H_2_O_2_ or O_2_·− are associated with increased ROS here. In Arabidopsis, there are several reports describing crosstalk between ROS and auxin (Hirt et al. 2000, Mangano et al. 2017, Huang et al. 2020, Blomster et al. 2011). Therefore, we cannot rule out the possibility that this small amount of ROS production, together with the reduction of auxin, contributed to the reduction in meristem size and promotion of root cell elongation in *S. parvula*. In *A. thaliana* seedlings, the same 100 mM NaCl treatment significantly suppressed root growth and caused a more than 10-fold increase in ROS throughout the root. This indicates severe salt damage caused by oxidative stress occurred.

Our proteomic analysis revealed several activations of antioxidant enzymes and responses. On the other hand, enzymes involved in protein translation and nucleic acid biosynthesis, which are essential for cell proliferation, decreased, consistent with suppression of DNA replication in meristems. More interestingly, among proteins related to amino acid metabolism, P5CDH level significantly increased. Proline is a well-known compatible solute that protects cellular membranes and enzymes in response to salt, cold, and drought stress (Alvarez et al. 2021). P5CDH is an enzyme that converts the toxic proline metabolism intermediate 1-pyrroline-5-carboxylate (P5C) to glutamate. Previously, it has been demonstrated that *S. parvula* accumulates proline in response to salt stress (Uzilday et al. 2015). However, the expressions of both anabolic genes (P5C synthetase and P5C reductase) and P5CDH were induced indicating enhanced proline cycling activity (Uzilday et al. 2015). This mechanism has been proposed to act as a redox shuttle between cytoplasm and mitochondria and a way to dissipate energy since NADPH is used for biosynthesis and FADH_2_ is produced during catabolism. We also found that GDH1 and GLN2 were significantly downregulated. This suggests that it may increase glutamate content without increasing proline content under moderate salinity. Increased glutamate might be converted to glutathione (Bosnan and Brosnan, 2013), which strengthens cellular defense systems and is also involved in direct signaling of abiotic and biotic stress responses (Qui et al. 2020).

Among proteins related to protein synthesis levels of translation initiation factor 5A-2 (FBR12) and 4A-1 (EIF4A1) and translation elongation factor TU and EF1B were decreased. It is well known that salinity has a profound effect on the regulation of protein biosynthesis as well as protein degradation (Hildebrandth, 2018). In a leaf proteomic study, Arabidopsis showed a contrasting response and salinity caused induction in levels of several translation initiation and elongation factors (eIF3A, eEF1B alpha 2 subunit, eEF2). However, similar to *S. parvula* such induction was not observed in leaves of halophyte *Thellungiella halophila* (Pang et al. 2010). Besides protein synthesis, turnover of proteins, hence degradation of some proteins for synthesis of new ones, is vital for plants to adapt changing environment. Among these proteins, regulatory particle triple-A ATPase 3 (RPT3), which is an ATPase subunit of 26S proteasome involved was induced. Similarly, a pyroglutamyl peptidase (peptidase C15 family cysteine protease) involved in proteolysis was also induced. On the other hand, 20S proteasome subunit PAB1 levels decreased. Among proteasome complexes, 26S proteasome is ubiquitin depended and selective while 20S proteasome is ubiquitin independent (Kurepa et al. 2009). Interestingly increase in 26S proteasome activity (over-expresser lines) has been linked to increased root elongation under drought in Arabidopsis and loss of 26S proteasome decreased primary root growth (Kurepa et al. 2009). Here we observe a similar phenomenon with moderate salinity induced primary root elongation in *S. parvula*. Another vital point regarding protein metabolism is proper folding of proteins, which is mediated by a plethora of chaperones, folding helpers and quality control mechanisms. However, chaperone proteins not only help fold proteins but regulate activity of signaling pathways. Among detected proteins, heat shock protein (HSP)70-HSP90 organizing protein HOP2, which has been shown to regulate salt tolerance by affecting the nucleo-cytoplasmic partitioning of HSP90 (Zhang et al., 2022) and MEC18.18, an activator of HSP90 was induced. Moreover, HSP90 has been shown to interact with TIR1 (auxin receptor) and control the distribution of PIN1 to regulate root growth in Arabidopsis (Samakovli et al., 2021). Therefore, HSP90 might be involved in primary root elongation in *S. parvula* via not only its chaperone activity but also its signal function through regulation of auxin signaling.

Proteomic analysis also shows that moderate salt stress changes carbon flux through glycolysis (phosphoglycerate mutase = PGM1, enolase = LOS2, fructose-biphosphate aldolase = FBA8). Among these, PGM1 and LOS2, enzymes that catalyze the last two steps of glycolysis decreased while FBA8 was induced. FBA8 catalyzes the reversible reaction of splitting fructose 1,6-bisphosphate into triose phosphates and is involved in both glycolysis and gluconeogenesis (transformation of non-carbohydrate substrates to glucose). These findings indicate that flux through glycolysis is decreased while gluconeogenesis is induced at moderate salinity. This hypothesis is also supported by lower level of pyruvate dehydrogenase E1 (MAB1), a subunit of pyruvate dehydrogenase complex, which links the glycolysis to the TCA cycle by conversion of pyruvate to acetyl-CoA. Induction of gluconeogenesis might support the synthesis of glucose-6-phosphate as a precursor for cellulose and cell-wall synthesis (Lan et al., 2013) required for primary root elongation at moderate salinity.

Two proteins related to cytoskeleton *ACT7* and *ACT8* levels decreased with moderate salinity. Among these *ACT8* belongs to vegetative subclass 1 actins, while *ACT7* belongs to vegetative subclass 2 actin and these two isovariants have diverse functions (Pei et al., 2012). In Arabidopsis, *ACT7* is responsible for initiation of root hairs and is regulated by auxin (Kannadasamy et al., 2001). On the other hand, *ACT8*, along with *ACT2* is responsible for root hair elongation. Decrease in levels of these proteins with moderate salinity could explain the observation of decreased density of root hairs and shorter root hair lengths.

Stress responsive proteins such as catalase (CAT) and GSTs (GSTU19, GSTF8 and GSTF9) might be linked to an adaptive response for regulation of cellular redox under moderate salinity and this is well-documented for salt stress (Verslues et al., 2007; Qi et al., 2010). Moreover, annexins (ANNAT7), LEAs (LEA26) or dehydrins (ERD10) are known to respond to various stress conditions including salt stress (Kosova et al., 2013). These proteins have diverse functions such as osmo-protection, increasing membrane stability and defense against ROS, which might be related to adaptation to increased salt concentration. Moreover, among the proteins identified, PYK10, encoding a β-glucosidase, is involved in the formation of ER body, which is a specific domain of the ER that is observed only in roots and seedlings of Brassicaceae species (Yamada et al. 2011). A previous proteomic analysis of the ER body of Arabidopsis demonstrated that PYK10 interacts with jacalin related lectins (JCLs,) and GDSL lipase proteins (Nagano et al., 2008). Interestingly, a JCL (MLP-300B) and a GDSL lipase (GGL18) was significantly decreased with moderate salinity in *S. parvula*, although PYK10 was induced. These findings suggest that moderate salinity regulates ER body protein profile in *S. parvula*. Previous studies demonstrate that PYK10 is involved in biotic stress tolerance through glucosinolates (Nakano et al., 2017), however further studies are required for elucidation of the involvement of ER body in response to salt stress and plant root growth regulation.

Here we found an important different response, primary root elongation under salinity conditions between *S. parvula* and *A. thaliana*. Recently, divergence in the abscisic acid (ABA) gene regulatory network has been reported in four Brassicaceae species (Sun et al., 2022). Application of exogenous ABA inhibits primary root elongation in all species except *S. parvula*. In Arabidopsis, ABA induces the expression of auxin biosynthetic genes such as *TAA1* and *YUC2*, and excess auxin inhibits root growth by suppressing cell elongation rate (Velasquez et al., 2016, Sun et al., 2022). Therefore, ABA-mediated regulation of auxin biosynthesis is thought to be lost in *S. parvula*. Moreover, primary roots of *S. parvula* are reversely elongated by ABA application (Sun et al., 2022). This was similar to our findings that cell elongation in elongation zones and suppression of meristematic tissue size occurred under moderate salinity where the auxin biosynthetic genes *TAA1* and *YUC8* were suppressed. Primary root elongation by suppression of auxin biosynthesis in *S. parvula* seedlings, together with the expression of stress-related proteins, may represent a novel strategy to reach deeper, less saline soils earlier.

## Acknowledgements

We would like to thank Shusei Sato and Masaru Bamba for their support in proteome analyses, Mika Teranishi and Kaoru Yoshiyama for teaching some experimental methods and Miki Otsuka for the technical support. This work was funded in part by JSPS KAKENHI grant number JP18H03947 (A.H). K.S obtained a scholarship from the Ministry of Education, Culture, Sports, Science, and Technology (MEXT).

## Author contributions

K. S, B. U., R. O, A. H. and I. T. conceived and designed the study. K. S, N. H., R. O., B. U., and A. H. conducted experiments and analyzed the data. K. S, B. U., A. H. and I. T. wrote the manuscript. All authors read and approved the final paper.

## Data Sharing and Data Availability

All data generated or analyzed during this study are included in this article and its supplementary information files.

## Supplemental Figure legends

**Supplementary Figure 1.**
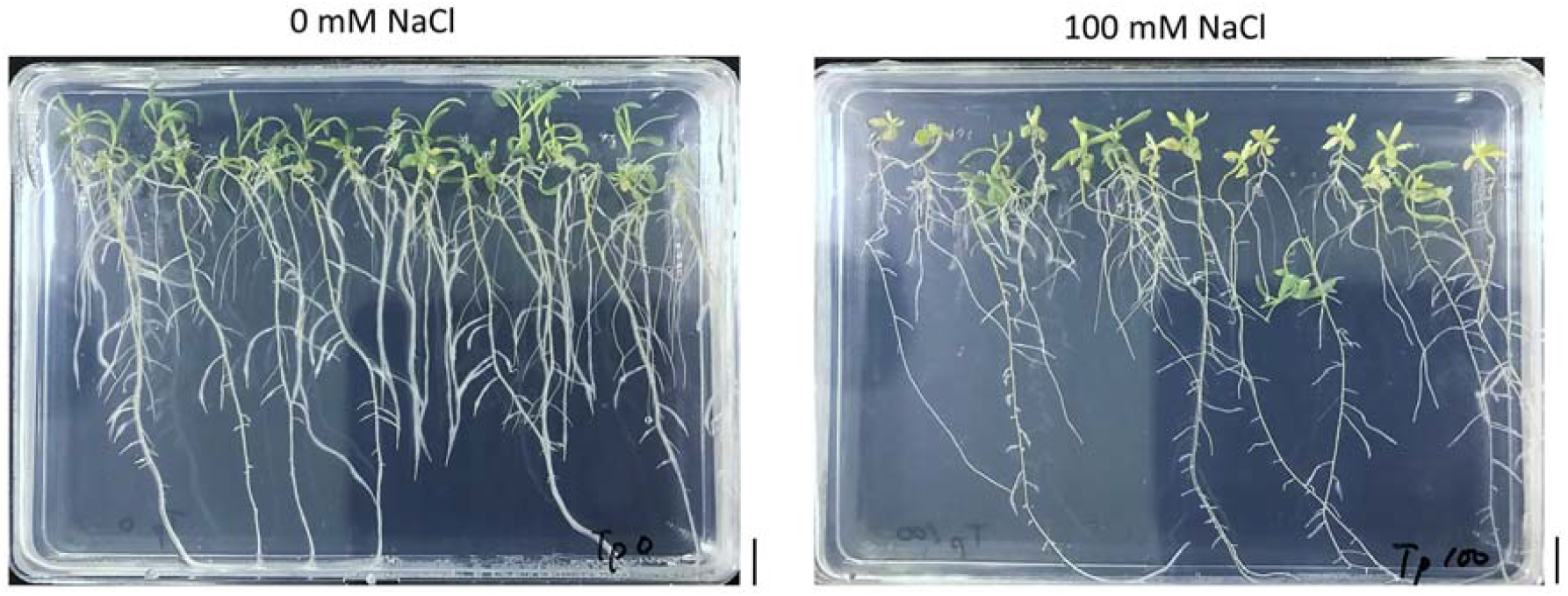
14-day-old *S. parvula* seedlings cultivated for 10 days on 1/2 MS sucrose agar containing 0 and 100 mM NaCl. Scale bar: 1 mm.

**Supplementary Figure 2.**
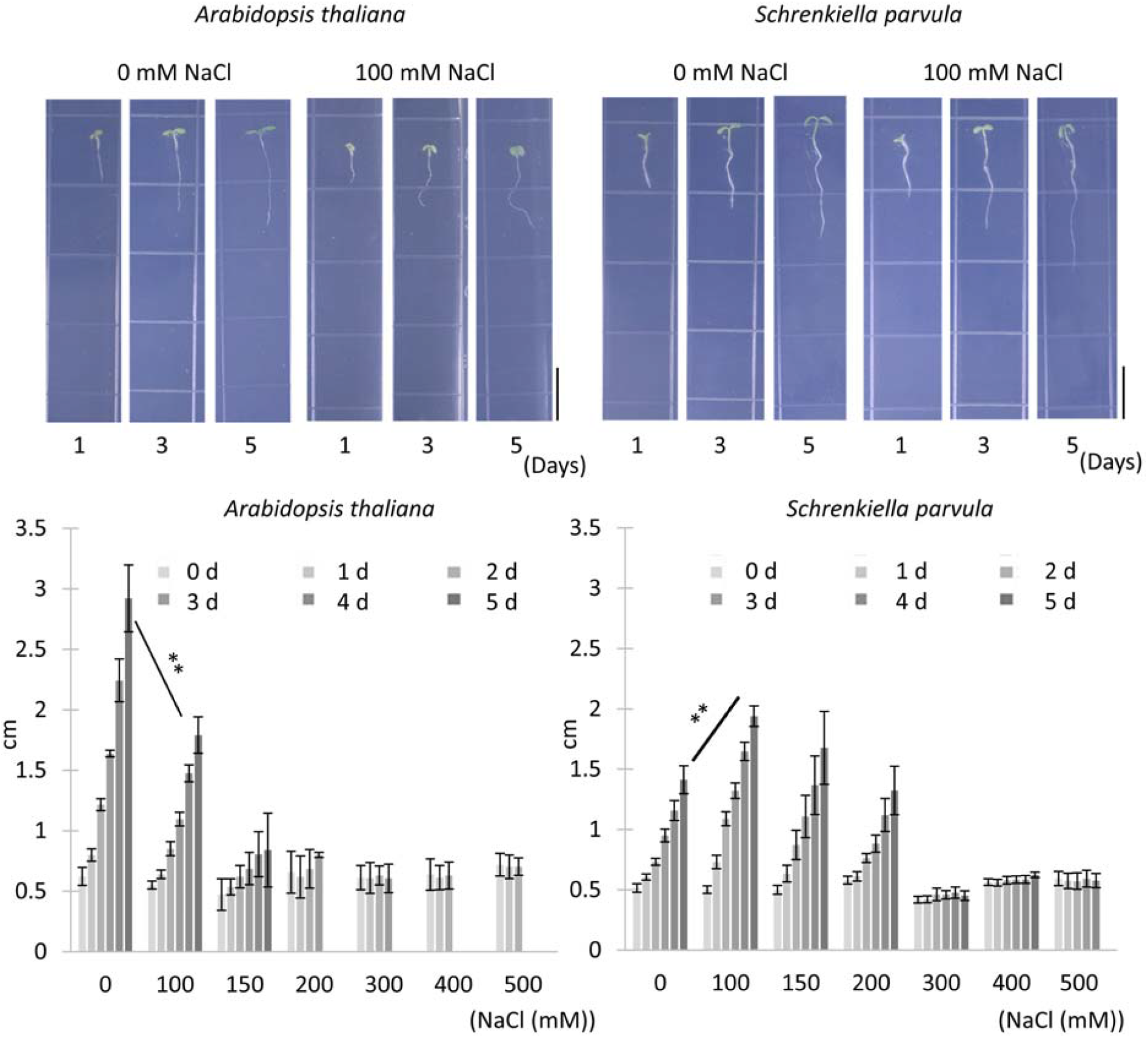
Growth of 0.5 cm root length seedlings of *A. thaliana* and *S. parvula* at under different NaCl concentrations for 5 days. Typical seedling images cultured in 0 and 100 mM NaCl for 1, 3 and 5 days. Scale bar: 1 cm. Root length was measured by Image J software. Data represent *n* = 20 < and means ± S.E. Student’s *t* test ** *p* < 0.01 between 0 and 100 mM NaCl in each plant for 5 days.

**Supplementary Figure 3.**
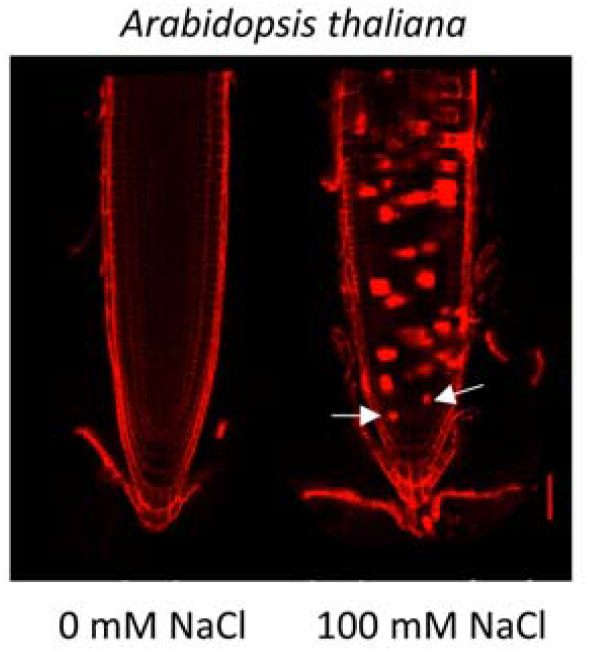
Confocal microscopy image of PI-stained root meristems of 6-day-old *A. thaliana* seedlings cultured in 0 mM and 100 mM NaCl for 2 days. White arrows indicate cell death in the apical meristem area. A further increase in these damaged dead cells was observed in *A. thaliana* compared to *S. parvula* (Fig. 4). Scale bar: 50 μm.

**Supplementary Figure 4.**
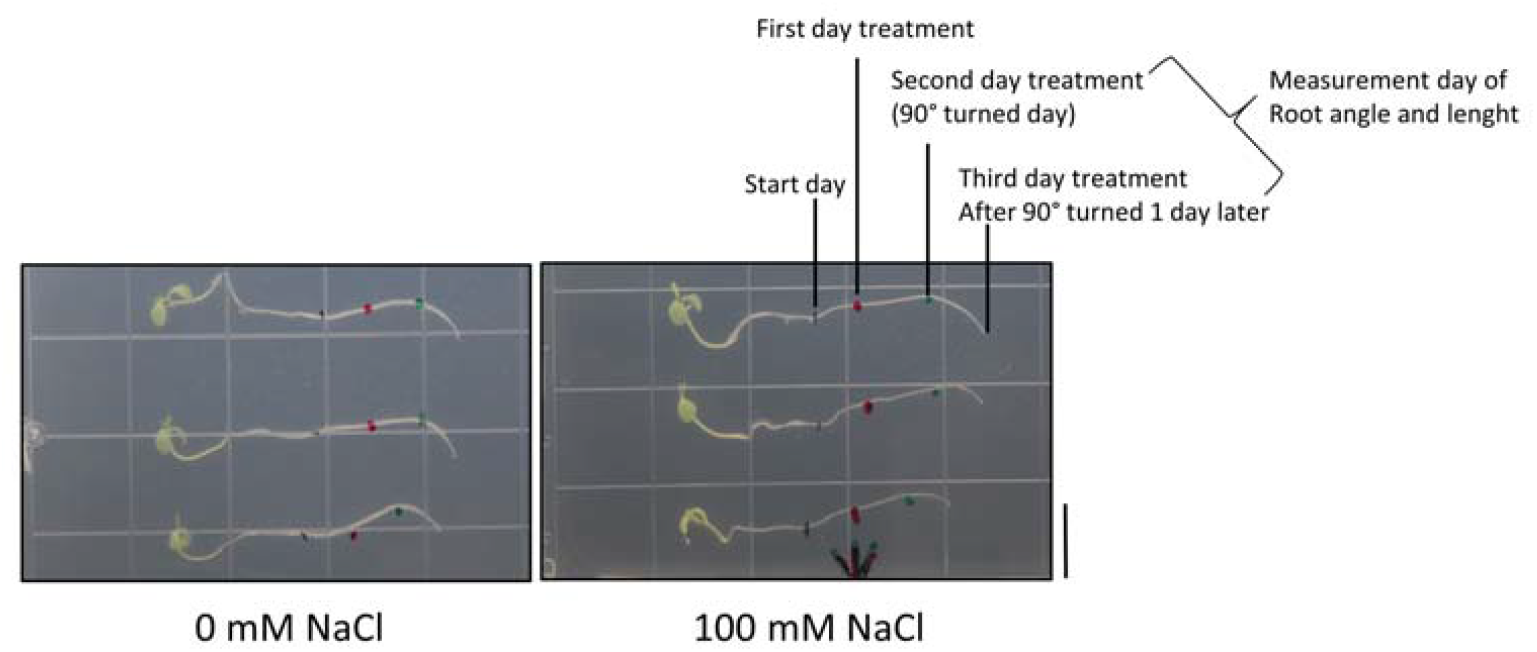
Effect of moderate salt on root gravitropic responses in *S. parvula* seedlings. Four-day-old seedlings were transferred to fresh medium with or without 100 mM NaCl and grown for 2 days before being tilted horizontally. After 90° rotation, images were taken after 24 hours. Scale bar: 1 cm.

**Supplementary Figure 5.**
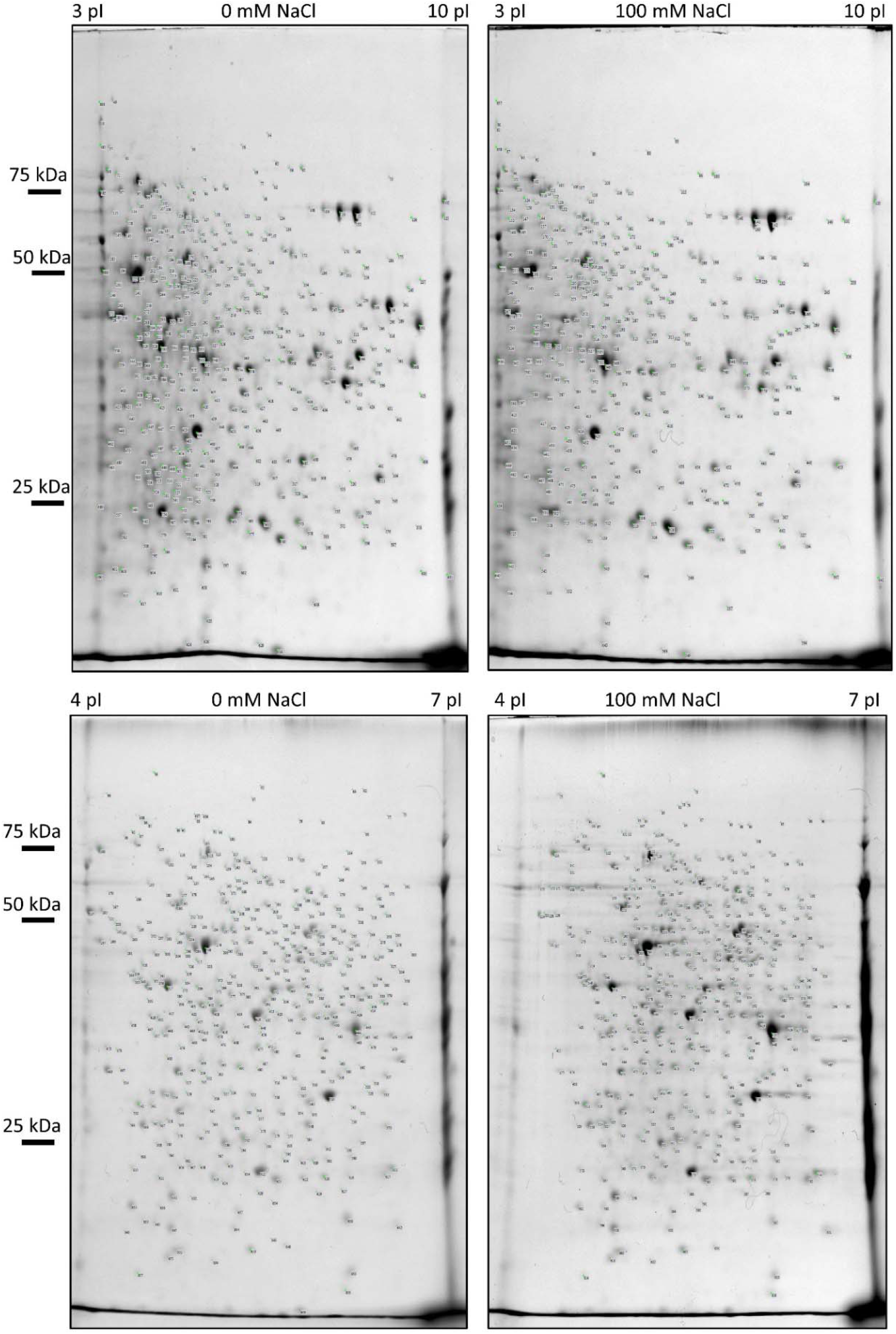
2DE images of total protein extracted from roots of 9-day-old *S. parvula* seedlings cultured in 0 mM and 100 mM NaCl for 5 days. pH 3-10 NL (upper panels) and pH 4-7 (bottom panels) IPG strips were used for first dimensional electrophoresis. Numbers indicate spot IDs of the identified proteins shown in Supplementary Tables 1 and 2.

